# 28 Minutes Later: Investigating the role of aflatrem-like compounds in Ophiocordyceps parasite manipulation of zombie ants

**DOI:** 10.1101/2022.09.08.507134

**Authors:** William C. Beckerson, Courtney Krider, Umar A. Mohammad, Charissa de Bekker

## Abstract

Coevolutionary relationships between parasites and their hosts can lead to the emergence of diverse phenotypes over time, as seen in *Ophiocordyceps* fungi that manipulate insect and arachnid behaviour to aid fungal spore transmission. The most conspicuous examples are found in ants of the Camponotini tribe, colloquially known as “zombie ants”. While the behaviours induced during infection are well described, their molecular underpinnings remain unknown. Recent genomics and transcriptomics analyses of *Ophiocordyceps camponoti-floridani* have identified several highly upregulated biomolecules produced by the fungus during infection of *Camponotus floridanus*. Among them is an ergot alkaloid related to the mycotoxin aflatrem, known to cause “staggers syndrome” in cows. Staggering, defined as unsteady movements side to side, is also observed in *C. floridanus* ants during late-stage infection. To test if aflatrem-like compounds could be responsible, we injected healthy ants with aflatrem and recorded their behaviour for 30 minutes. Using both the automated object-tracking software MARGO and manual behavioural quantification, we found that aflatrem reduced ant activity and speed, and increased staggering behaviours. To examine underlying transcriptomic changes, we performed RNA-Seq on the heads of aflatrem-injected ants, keeping in step with previous transcriptomic work on *Ophiocordyceps*-manipulated ants. We identified 261 genes that were significantly dysregulated in the aflatrem-injected ants compared to sham-injected controls. When compared with RNA-Seq data from *Ophiocordyceps*-manipulated ants, we found that both groups shared 113 differentially regulated genes. These included *sensory neuron membrane protein* genes, several *odorant-binding protein* genes, and musculoskeletal genes such as *titin* and *obscurin*. Together, these results indicate that aflatrem-like compounds significantly affect neuromuscular and sensory function in *C. floridanus*. The conservation of staggers phenotype between *C. floridanus* and *Bos taurus* suggests that behaviour manipulating strategies exhibited across the Tree of Life may be more similar in approach, if not widely different in application, than we realize.

## 1. INTRODUCTION

When a parasite infects a host, an intense co-evolutionary relationship can develop over time, driven by selective pressures exhibited by the host acting as the new environment for the parasite. As the parasite becomes more successful, the selective burdens shift to favour emergent resistance phenotypes in the host. Over time, these reciprocal selective pressures can lead to host-specificity and the subsequent emergence of unique manipulation phenotypes (Fain 1994; Flor 1956; McLaughlin & Malik, 2017). Host manipulations provide an adaptive advantage to the parasite, for instance through the induction of specific behaviours that increase its reproduction success (Sánches and Biron, 2019). These adaptive phenotypes are different from sickness behaviours, defined as actions exhibited by infected individuals that aid in the fitness of the host (e.g., lethargy) (Breed & Moore, 2010), and non-adaptive behaviours that benefit neither the host nor the parasite. One clear example of behavioural manipulation can be observed in the parasitic flat worm *Leucochloridium paradoxum*, which manipulates amber snails from the genus *Succinea*. Once inside a snail, the worm drives the host to ascend nearby structures, a common manipulation behaviour known as “summit disease” (Wesołowska and Wesołowski, 2013). Summiting behaviour makes snails more susceptible to predatory birds, the primary host of *L. paradoxum*, ultimately increasing the parasite’s likelihood of completing its lifecycle (Wesołowska and Wesołowski, 2013).

Such behaviour-manipulating parasites are often colloquially referred to as “zombie parasites”. Not because they reanimate the dead, but because the drastically altered behavioural phenotypes exhibited by infected hosts favour parasite transmission, which is reminiscent of the behaviours of zombies in pop culture and sci-fi media (e.g., the movies *28 Days Later* and *World War Z*). While the manifestations of behavioural manipulation vary across species, many host-manipulating parasites have convergently evolved to induce summiting behaviour to increase their transmission (Araújo & Hughes, 2019; de Bekker et al., 2021; Latchininsky et al., 2016; Steinkraus et al., 2017; Wesołowski & Wesołowski, 2013). One of the largest known groups of parasites that cause summit disease are the “zombie fungi” of the genus *Ophiocordyceps*, which infect various species of insects and arachnids globally (Arruda et al., 2021; Cooley et al., 2018; de Bekker et al., 2021). This includes wasps (e.g., *O. humberti* (Sobczak et al., 2020)), beetles (e.g., *O. curculionum* (Shrestha et al., 2016)), scale bugs (e.g., *Ophiocordyceps clavulata* (Petch, 1933)), caterpillars (e.g., *O. sinensis* (Guo et al., 2021)), flies (e.g., *O. dipterigena* (Chhetri et al., 2020)), and spiders (e.g., *O. engleriana* (Sung et al., 2007)). However, a large portion of the currently described species belong within the species complex *Ophiocordyceps unilateralis*; one of the more well-studied groups of zombie fungi that collectively infect and manipulate carpenter ants with extreme host-specificity (Araújo et al., 2018, Araújo et al., 2020).

*Ophiocordyceps unilateralis* species begin their lifecycle as a spore, ejected from their parental fruiting body. These spores sail on the wind to infect new ants, either through immediate contact or through the production and attachment of secondary capillispores from the forest floor (Evans et al., 2018). Once inside the ant, the fungus begins to secrete an array of biomolecules in a time-specific manner; some of which alter the behaviour of their host (de Bekker et al., 2015; Will et al., 2020). The first notable change occurs as these ordinarily social hosts begin to show a lessened ability to communicate and participate in foraging tasks (Trinh et al., 2021). Subsequently, infected ants abandon the nest and climb nearby vegetation. Once at an elevated position, they latch on with their mandibles, such that the ant is perched on the underside of its biting substrate (Andersen et al., 2009). The fungus then completes its lifecycle through the formation of an endosclerotium that siphons the remaining nutrients from the host for the formation of a new fruiting body (Fredericksen et al., 2017). Closer observation and quantification of behaviours in both the lab and the field have led to the identification of other, more subtle behavioural changes. Altered behaviours include changes in daily rhythms (Hughes et al., 2011; Trinh et al., 2021) and the loss of motor coordination and balance, resulting in ants that fumble their way around as in a “drunkard’s walk” (Supplemental Video 1) (Hughes et al., 2011; Trinh et al., 2021; Will et al., 2020). These conspicuous behavioural changes caused by the fungus during infection make the *O. unilateralis* species complex and their hosts particularly well-suited for the study of behaviour-manipulating parasites in greater detail (de Bekker et al., 2015).

Recent work establishing fungal culturing and ant infection techniques, as well as the genomic sequencing of several *O. unilateralis s.l.* species, has laid the groundwork for research aimed at understanding the molecular framework driving behavioural manipulation (de Bekker et al., 2015; de Bekker et al., 2017; Will et al., 2020). Transcriptomics analyses of the North American species *Ophiocordyceps kimflemingiae* and *Ophiocordyceps camponoti-floridani* have identified secreted proteins and secondary metabolite clusters highly upregulated during summiting behaviour (de Bekker et al., 2015; Will et al., 2020). One of these secondary metabolite clusters demonstrated a ∼5,900 (in *O. kimflemingiae*) and ∼12,000-fold (in *O. camponoti-floridani*) increase in expression during manipulation compared to fungi grown in culture (Will et al., 2020). This cluster was first identified as an ergot alkaloid-producing pathway based on the functional annotation of its tryptophan dimethyltransferase (Pfam: TRP_DMAT (PF11991)) backbone gene (de Bekker et al., 2015). More detailed annotation of the genes within and surrounding the cluster demonstrated that it was homologous to the aflatrem-producing cluster in the plant pathogen *Aspergillus flavus* (Will et al., 2020). Aflatrem, a secondary metabolite of the indole-diterpene group called aflatoxins, is a potent tremorgenic and carcinogenic mycotoxin known to cause neurological disorders in animals (Duran et al., 2006; Nicholson et al., 2009). Homologs to the atmQ, atmD, atmP, atmM, atmC, atmB, atmG, and idtS genes involved in the synthesis of aflatrem in *A. flavus* were all found in both the *O. camponoti-floridani* and *O. kimflemingiae* genomes (Will et al., 2020). Further genomic comparisons with other *Ophiocordyceps* genomes revealed that this secondary metabolite cluster appears to be conserved across species within *O. unilateralis* but not in other *Ophiocordyceps* species outside the species complex (e.g., *Ophiocordycpes australis s.l.*) (de Bekker et al., 2017). Taken together, this suggests that *Ophiocordyceps* species that infect ants of the tribe Camponotini can produce a secondary metabolite that is relatively similar to aflatrem, which they upregulate at the time when ant behaviour is being manipulated. This begs the question; what role might aflatrem-like compounds play in the manipulation of *Camponotus* behaviour?

The effects of aflatrem on animal behaviour have previously been studied in *Bos taurus* where the symptoms of aflatrem exposure were first observed in livestock that unintentionally ingested *A. flavus*-contaminated grain feed (Selala et al., 2008; Valdes et al., 1985). In these cases, cows exhibit neurological problems exhibited by muscle tremors and hyperexcitability, followed shortly by incoordination, ataxia, and seizures (Selala et al., 2008; Valdes et al., 1985). These neurological symptoms, linked directly to aflatrem’s positive allosteric modulation of Gamma-aminobutyric acid (GABA) receptors, are referred to as “staggers syndrome” (Eldefrawi et al, 1990; Grant et al., 1987; Yao et al., 1989). Should aflatrem-like compounds exhibit similar effects in *Camponotus* species, it could have important ramifications for host manipulation given the suspected involvement of GABA receptors in caste and behavioural differentiation in *Camponotus* ants (Gospocic et al., 2017; Graham, 2018). The likelihood that an aflatrem-like compound may be employed during the manipulation of carpenter ants is further supported by the “drunkard’s walk” observed during late-stage infection; a behaviour reminiscent of the staggers syndrome detailed in vertebrates (Supplemental Video 1). We, therefore, hypothesize that aflatrem-like compounds are responsible for the stagger phenotype observed during the behavioural manipulation of carpenter ants by *O. unilateralis* s.l. To test this hypothesis, we formulated two research questions: 1) Does aflatrem have any behavioral effect on carpenter ants and 2) how are genes dysregulated during induced behaviours? If aflatrem-like compounds are responsible for some of the motor impairment previously described as the “drunkard’s walk” in *Ophiocordyceps*-infected ants, then increasing doses of aflatrem should increase the duration and severity of staggering symptoms as previously studied in *B. taurus*. Furthermore, we expect to see that neuromuscular genes required for coordination should be dysregulated.

In this study, we administered aflatrem to the Florida carpenter ant *Camponotus floridanus*, the host of *O. camponoti-floridani* (from here simply referred to as *Ophiocordyceps*). We used microinjections to mimic the *in vivo* secretion of aflatrem-like compounds by fungal cells in the haemolymph during infection. Given the ease of its procurement, and its previous use in other animal models, we used high purity synthesized aflatrem (>98%) in our experiments (TePaske et al., 1992; Trienens & Rohlfs, 2011). Following injections, we observed and quantified ant behaviour to address if aflatrem can induce behaviours consistent with those exhibited during infection by *Ophiocordyceps*. Furthermore, to investigate which host pathways are affected by aflatrem and potentially give rise to the exhibited behavioural phenotypes, we also conducted transcriptomics analyses of aflatrem-injected individuals. Comparison of these results to transcriptomics data from *Ophiocordyceps*-infected *C. floridanus* ants (Will et al., 2020) yielded several consistent differentially expressed genes, including genes that encode neuromuscular and chemosensory proteins.

## 2. MATERIALS AND METHODS

### 2.1 **Ant Collection**

For use in behavioural and transcriptomics experiments, we collected three separate queenless *Camponotus floridanus* colonies of unknown age using a DeWALT Cordless 2 Gallon Wet/Dry ShopVac at the University of Central Florida’s arboretum, coordinates: Colony 1 (28.602748, - 81.194269), Colony 2 (28.603918, -81.191682), and Colony 3 (28.601430, -81.191251) (KML File 1). All colonies were collected from fallen logs using a tarp to ensure that nearly all ants that resided within were collected. In total, each collected colony contained between an approximated 100-200 major and minor worker ants without larvae, indicating that all were likely satellite nests of bigger colonies in the area. We housed these colonies in 9.4 L plastic containers (Rubbermaid) lined with talcum (Acros Organics), applied using a viscous 20% w/v solution of powder suspended in 100% ethanol to prevent ants from climbing the walls. Throughout the study, we provided the ants with fresh supplies of both double distilled water (ddH2O) and 15% sucrose solutions, *ad libitum*, as well as aluminium foil-wrapped falcon tubes that functioned as darkened nest chambers. We also kept the housing in an incubation chamber (Percival) set to the following program: 20°C – 0 lux – 80% relative humidity (RH) with a linear increase in temperature, light, and decrease in humidity over 4 hours to reach 28°C – 2,100 lux – 65% RH, at which conditions were held for 4 hours followed by a linear decrease in the temperature, light, and increase in humidity, returning to 20°C – 0 lux – 80% RH over 4 hours; and held for 12 hours. Only minor worker ants were used in this study, without formally determining their caste differentiation into foragers and nurses. However, the ants chosen for injection were outside and away from the darkened nest tubes at the time of collection, exploring the edges of the containers.

### 2.2 Microinjections

To test our hypothesis and identify the effect of aflatrem-like compounds on *C. floridanus*, we established a behavioural assay using microinjections to introduce aflatrem in a manner biologically relevant to the secretion of aflatrem-like compounds by *Ophiocordyceps* species in the haemolymph of their ant hosts (Figure 1).

**Figure 1.**
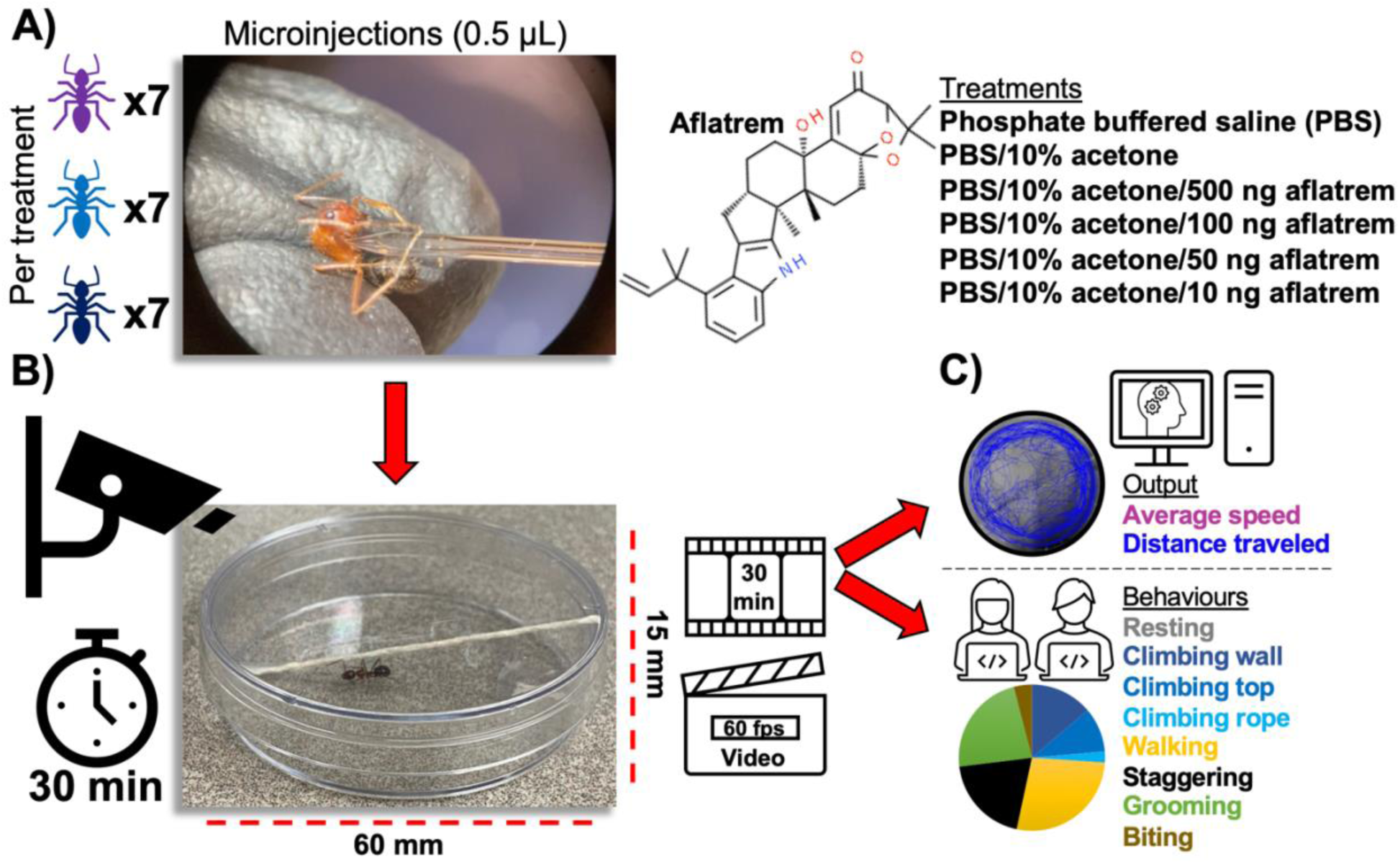
Experimental overview of (A) the treatment groups, each consisting of seven individuals from three different colonies of *Camponotus floridanus*, and an example of microinjection, (B) the arena setup for behavioural assays and parameters for the observational periods, and (C) the behavioural analyses using automated and manual behaviour scoring software.

For these assays, we utilized high purity aflatrem, >98% C_32_H_39_NO_4_, manufactured from *A. flavus* by BioCrick Science Solution Specialists in non-water-soluble crystal form. Without an approximation for the level of aflatrem-like compounds secreted in the host *in vivo*, we suspended the aflatrem to a concentration of 10 mg/mL in 100% acetone (Fisher Chemical). This allowed us to make further dilutions to reach concentrations informed by other studies on the effects of toxic compounds in insects (Buczkowski & Woosler, 2019; Klotz & Moss, 1996; Sakamoto & Goka, 2021). We performed injections using borosilicate glass micropipettes (Fisher) pulled with a glass puller (model: PC-100, Narishige) under the following parameters: single step, all weights, heated to 62°C. To inject the ants, we held each one by the dorsal side of the thorax before piercing the carapace on the ventral side between the first and second pair of legs, followed by slowly injecting 0.5 μL of solution into the ant while simultaneously releasing grip to alleviate intracavitary pressure. This procedure is similar to those routinely used to infect carpenter ants with fungal cells (de Bekker et al., 2014; de Bekker et al., 2015; Will et al., 2020; Trinh et al., 2021). Performing injections in the soft tissue between the joints allowed us to avoid puncturing the hard exoskeleton. This combined with constant pressure from the legs during walking holds the wound closed and prevents the ant from bleeding and allows them to recover.

Because the aflatrem used in this study is non-water soluble, we performed a preliminary trial to identify acetone tolerance in ants. As such, we injected subjects with 100%, 50%, 25%, 10%, and 5% acetone solutions diluted with phosphate buffered saline (PBS) (137 mmol NaCl, 2.7 mmol KCl, 10 mmol Na_2_HPO_4_, 1.8 mmol KH_2_PO_4_, pH adjusted to 7.4), along with a 100% PBS solution as a control. Ten minutes prior to injection, we removed containers housing ant colonies from the incubator and set them on the lab bench top to acclimate to lab conditions. We also performed all injections in triplicate, one ant per colony per treatment. After injection, we placed the ants into individual 60 mm x 15 mm petri dishes (USA Scientific) and returned them to the incubator. We kept these ants under observation for 48 hours post injection using an infrared GoPro Hero6 set to record time-lapse footage with 60 second intervals and a CMVision IR30 Illuminator lamp for observation during the night cycle. From these videos, we were able to easily differentiate dead and resting *C. floridanus* ants from one another; dying ants coiled their legs and fell to one side upon death while resting ants remained prone with legs supporting their bodies. As such, we were able to estimate the time of death of each ant within an interval of 1 minute by noting the frame in which they exhibited “death behaviours” (Supplemental video 2). We euthanized any ants that survived the 48-hour test period by freezing at -20°C for 24 hours. Results from these preliminary analyses indicated that a 10% acetone solution was the highest concentration for which there were no statistical differences in average survivability compared to PBS-injected and non-injected controls.

To test the survivability of ants post injection, and determine if any observed behavioural phenotypes may be attributed to “dying behaviours” resulting from aflatrem exposure, we performed a second preliminary trial. We injected 7 ants with 0.5 μL 10% acetone solution containing 500 ng aflatrem. This was the highest dose possible using the 10 mg/mL aflatrem stock. After injection, we recorded the ants for 48 h using the same GoPro setup as the previous preliminary test. We then compared the two groups and found no significant differences in mortality, with all individuals surviving for at least 24 h and an average time of death arising at 34 h. The results indicate that all ants that died in the trial did so as a result of desiccation from lack of water.

Confident that neither 10% acetone nor 500 ng aflatrem killed the ants or induced death-related behaviours, we conducted an assay to determine the effects of aflatrem on ant behaviour using 10% acetone solutions containing the following doses of aflatrem; 10 ng, 50 ng, 100 ng, 500 ng. For each dose of aflatrem, we injected 7 ants from each of the 3 colonies (N=21, per dose) for a total sample size of N=126. We individually lifted each ant from their housing by hand wearing neoprene gloves (NeoTouch) and allowed them to acclimate to the lab conditions by placing them in individual 60 mm x 15 mm petri dishes (USA Scientific) on the lab benchtop for 10 minutes. We fitted each petri dish with a thread of twine spanning the dish from the bottom of one side to the top of the other. This provided ants with a substrate for biting and more dexterous climbing behaviours (Figure 1-B). To allow ants to walk around the perimeter of the dish unimpeded, we made sure to tape (Scotch) the twine pressed flat against the bottom corner of the petri dishes. We created a new petri dish arena for each test subject (N=126) to avoid any behavioural changes that may be induced via chemical ques left by prior occupants. After the 10-minute acclimation period, we injected ants with aflatrem following the microinjection protocols established in the preliminary trials. The doses of aflatrem used were based on approximate ant equivalent weights, verified by measuring *C. floridanus* body volume through length and weight following each experiment. While we measured body lengths to the nearest 0.5 mm for each individual, weight was measured in groups of seven ants corresponding to their treatment and colony ID. We recorded these measurements immediately after the ants were euthanatized via freezing, preserving the approximate water weight for each ant. To account for the approximate weight of solution injected into each ant, we also subtracted 3.5 mg from the total weight of each group. Based on an average of 8.28 mg weight and 6.68 mm length calculated from measurements of all 126 subjects used in this study (Supplemental Table 1), the four doses of aflatrem that were used are approximately 6.04e-5, 1.21e-5, 0.60e-5, and 0.12e-5 ant equivalents. These ant equivalent doses are consistent with pervious works testing insecticidal compounds in ants (Buczkowski & Woosler, 2019; Klotz & Moss, 1996; Sakamoto & Goka, 2021).

Using an independent two sample t-test to compare the behaviour of ants injected with PBS to those injected with 10% acetone, we were able to rule out any significant differences between the two groups: rest (*p = 0.787*), speed (*p = 0.997*), distance (*p = 0.947*), stagger (*p = 0.205*). Even though neither group was exposed to aflatrem, we unexpectedly observed a few instances of staggering. To determine if a small amount of staggering behaviour is normal in ants, which would need to be accounted for in our results, we reviewed the footage at timestamps logged for these events with an increased level of scrutiny. Upon closer review, we discerned that nearly all staggering observed in both groups fell into the “gaster wagging” sub-type (Supplemental Video 3), and nearly always occurred in the first 30 seconds of the recording. This trend was also observed in the aflatrem treatment groups as well, indicating that this particular type of staggering behaviour is likely a transient physical response to the trauma resulting from the injection process. We confirmed this hypothesis through a quick follow-up experiment pricking ants with empty glass capillaries, which also induced gaster wagging in some.

### 2.3 Video scoring of behavioural assays post injection

To record behaviours post injections, we used an iPhone 13 pro. We recorded three or four arenas at a time in a rolling format to ensure high quality resolution at 60 fps. This generated video files approximately 45 minutes in length. In cases where ants exhibited signs of physical injury from the injection, e.g., immobility in the hind legs or body curling, we euthanized the subject and replaced it with a new ant and new arena. If any ants were exhibiting resting behaviour at the end of their recording period, we tapped their arenas several times to induce movement. All instances of tapping elicited activity, confirming that all test subjects were still alive.

To crop the rolling videos into clips showing only individual arenas, we first converted the iPhone .MOV files to .mp4 files using iMovie. We then exported these .mp4 files to Movavi Video Editor Plus 2022 and trimmed each clip to exactly 30 minutes, beginning as soon as each arena was positioned in the field of view. The names for each of these clips were blinded before two observers manually scored the videos using the behaviour tracking software CowLog3 (Pastell, 2016). Before formal scoring, we trained each observer using eight test run videos; four of ants injected with 500 ng aflatrem, and four of ants injected with PBS. These videos were made during proof-of-principle testing and are not included in the data presented in this study. We used the results from these practice runs to form a consensus on scoring methodology. From these training videos, the observers identified eight scorable behaviours: Climbing Wall, Climbing Top, Climbing Rope, Walking, Staggering, Grooming, Resting, and Biting (Supplemental Video 3). While most of these behaviours are self-explanatory, we defined “staggering” as a slow, sluggish walking behaviour with repeated lateral movements (Supplemental Video 4). We also categorized behaviours as “staggering” when trouble balancing, listing to one side, falling over, walking in tight circles, flipping over, or dramatic wagging of the gaster were observed (Supplemental Video 4). Additionally, each observer kept notes on unique behaviours observed during the scoring period, including walking backwards, lunging/bucking, and two instances of convulsions (Supplemental Video 5).

During the formal scoring, we assigned the blinded videos to the observers in a randomized block format (Supplemental Figure 1-A) with each observer scoring three videos at random per treatment group per colony. Both observers scored the seventh video for each treatment group, from which the results were compared and used as a formative metric to ensure consistent scoring throughout the evaluation period. While most behaviours were readily distinguishable, resulting in very similar scores between observers, walking and staggering provided the most discrepancy. We, therefore, used staggering as the primary measurement to evaluate scoring consistency across observers (Supplemental Figure 1-B).

In addition to manual observations, we also analysed each video using the MATLAB-based automated object-tracking software Massively Automated Real-time GUI for Object-tracking (MARGO) (Werkhoven et al., 2019). Using this tool, we were able to measure the speed and distance travelled by each subject. While we calculated distance travelled by measuring the total distance between centroid coordinates over the entire 30 min duration, we only measured speed, calculated in mms travelled per second, during activity. To optimize the tracking within the field of view for each video, we used the following metrics: region of interest (ROI) was set to circular shape and manually overlayed around the arena, background referencing was set to detect dark objects on light backgrounds, tracking threshold was set between 70-80, distance of the arena was set to 60 mm using the custom measuring tool, frame rate was set to 30, and min/max area was set to 5 and 80, respectively.

Before testing the effects of aflatrem on ant behaviour, we compared the results from the 10% acetone controls to the 100% PBS controls to identify any potential behavioural phenotypes induced by the addition of acetone to the injections. To test for any statistical significance between the two groups, we used a two-sample t-test in RStudio v1.3.959. Following the comparison of these two controls, we tested the effect of aflatrem exposure on ant behaviour with a linear regression model using the lme4 package in RStudio, by comparing the 10% acetone control (0 ng aflatrem) with 10% acetone containing 10 ng, 50 ng, 100 ng, and 500 ng aflatrem treatments.

### 2.4 RNA sequencing

To understand the effects of aflatrem exposure in *C. floridanus* at the genomic level, we injected a total of 10 worker ants from colony 1 with either 10% acetone (control group, N=5) or 500 ng aflatrem (treatment group, N=5). We then placed each ant into their own petri dish arena for 4 minutes. We chose this timepoint based on the scaling levels of activity observed in our behavioural assays (Supplemental Figure 2). After this period, we quickly placed the ants into 2 mL microcentrifuge tubes and snap froze them in liquid nitrogen. Subsequently, we removed the head from each ant with forceps and placed it in a pre-frozen 2.0 mL Self-Standing Impact Resistance Microtube (USA Scientific) containing 2 Grade 25 5/32” metal ball bearings (Wheels Manufacturing) for mechanical tissue disruption in a SPEX® SamplePrep 1600 MiniG at 1500 rpm for 30 sec. We dissolved the frozen samples in 500 uL TRIzol (Ambion) to extract the RNA and isolated it using the Qiagen RNeasy® MiniElute® Cleanup Kit (Will et al., 2020). The resulting RNA was stored at -80C and shipped to Azenta US for Next Generation Sequencing. Libraries were prepared by Azenta US using PolyA selection and sequenced as 150 bp paired end reads on an Illumina HiSeq, resulting in ∼350M reads. Read quality was verified using FastQC (Andrews 2010). We used Trimmomatic (Bolger et al., 2014) to trim reads for adapters and quality using the following parameters: Leading: 3, Trailing: 3, MinLen: 36. We then aligned the transcripts to the *Camponotus floridanus* reference genome (Gene bank ascension number GCA_003227725.1) using Spliced Transcripts Alignment to a Reference (STAR) 2.7.10a (Dobin et al., 2013). After alignment, we checked the resulting .bam files for quality using the standard metric of uniquely mapped reads > 70%, sorted, and indexed using Samtools (Danecek et al., 2021). We next calculated the number of gene hits for each sample using FeatureCounts, following the suggested counting of strictly uniquely mapped reads within exonic regions (Liao et al., 2019). The quality of the resulting files was validated using a benchmark of 15-20 million reads per sample. After removing hits with low level of expression (counts less than 100 per gene, i.e., an average less than 10), we normalized the remaining counts using DESeq2 (Love et al., 2014) eliminating 3,227 genes from further analysis. We performed differential expression analysis on the remaining 10,443 genes using DESeq2 with an alpha value of 0.05 and functionally annotated the differentially expressed genes using Blast2GO (Conesa et al., 2005). To visualize the overlap between the differentially expressed genes found in this study and those found in a previous transcriptomics dataset obtained from *Ophiocordyceps*-infected ants (Will et al., 2020), we developed a Kaleidoscope Diagram using a custom R package made by Andrew Swafford (Supplemental R-package 1). Finally, we performed a GO enrichment analysis using the R package from Das & de Bekker (2022) made publicly available at https://github.com/biplabendu/timecourseRnaseq.

### 2.5 Ethical Note

We performed all injections and behavioural analyses in this study on the invertebrate ant species *Camponotus floridanus*, which is not endangered and is a prevalent species in Central Florida. We collected colonies of this species at the University of Central Florida Arboretum, which did not require prior approval from an animal welfare or ethics committee. Only three colonies were collected to meet the minimal scientific standard of triplicate replication. Collection utilized a fast and handling-minimizing method. We applied the following standards in effort to reduce the amount of stress incurred by the ants throughout our study as much as possible. Housing was conducted in an environmental chamber set to mirror the subjects’ natural conditions. We maintained colonies by providing *ad libitum* food and water, and dark enclosures for nesting. We furthermore conducted experiments on the minimum required number of subjects to reach the sample size needed for statistical analysis. Ants were quickly euthanized at -20°C in instances where they were injured during treatment and immediately following all experiments.

## 3. RESULTS

### 3.1 The behavioural effects of aflatrem in *C. floridanus*

Confident that behaviours were not significantly affected by the addition of 10% acetone in the injections, we made comparisons across increasing doses of aflatrem. A linear regression model showed that time spent resting in the arena was positively correlated with increasing doses of aflatrem (*F*_4,105_ = 5.54, *p* = 2.05e-2) (Figure 2-A), while the average speed of aflatrem-treated ants was negatively correlated (*F*_4,105_ = 4.91, *p* =2.89e-2) (Figure 2-B). However, the average distance travelled did not correlate with aflatrem treatment (*F*_4,105_ = 3.036, *p* = 8.44e-2, ns) (Figure 2-C). Finally, bouts of staggering did positively correlate with increasing doses of aflatrem (*F*_4,105_ = 36.65, *p* = 2.32e-8) (Figure 2-D).

**Figure 2.**
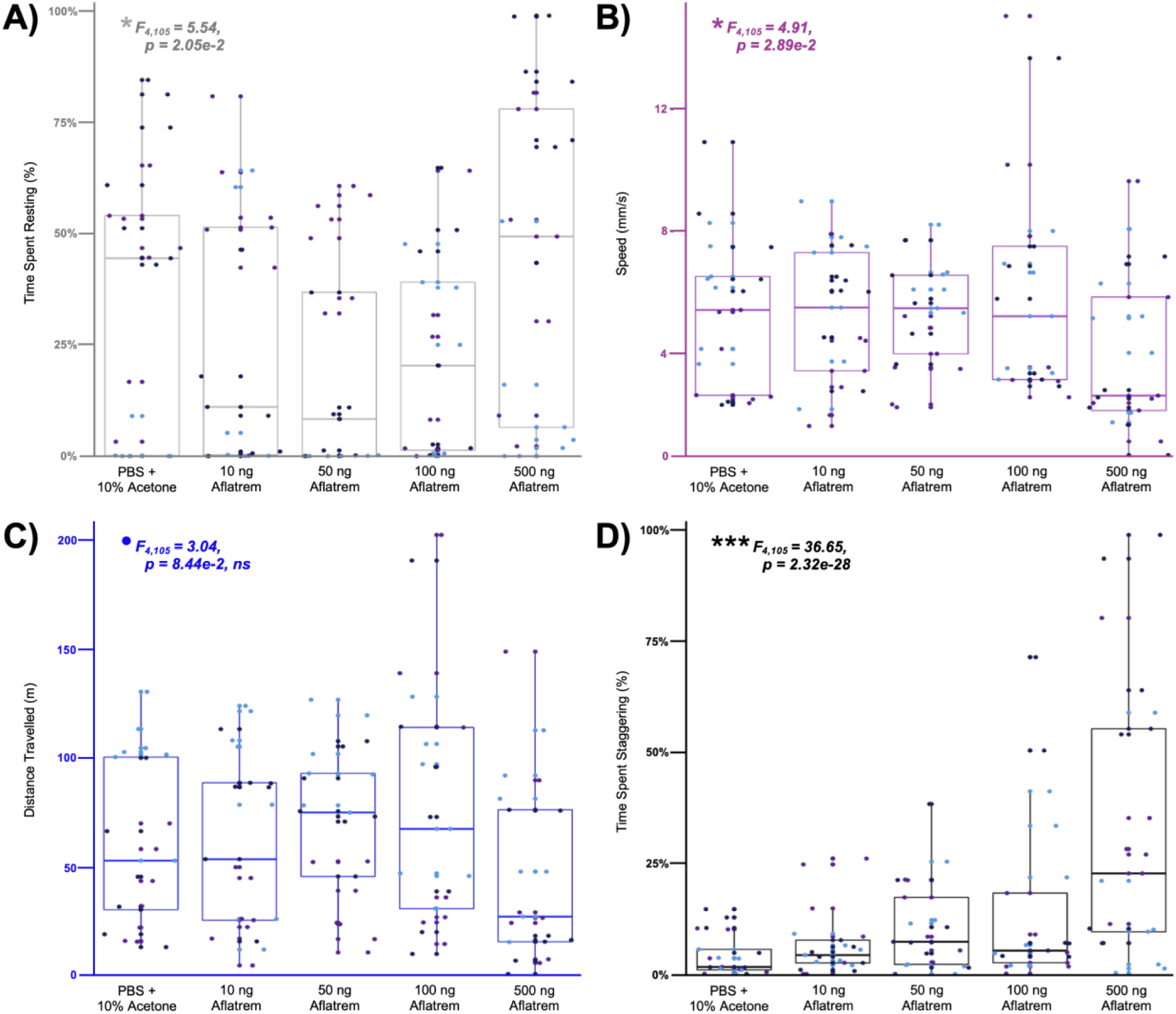
Behavioural results collected from *C. floridanus* ants injected with increasing doses of aflatrem. Each boxplot depicts the quartiles of the distribution for each treatment with ants from colonies 1, 2, and 3 shown as purple, light blue, and dark blue dots, respectively. A) shows the proportion of time spent resting for each ant, B) shows their average speed during activity in millimetres per second, C) shows their total distance travelled in meters, and D) shows the proportion of activity spent staggering for each ant. The linear regression model results are shown in the top left of the graph for each behaviour.

### 3.2 The effects of aflatrem on gene expression in *C. floridanus*

While injection of *C. floridanus* with aflatrem resulted in quantifiable phenotypes, we next asked which genes could be underlying the emergence of staggering and diminished activity phenotypes. As such, we conducted transcriptomics to compare gene expression in the heads of ants treated with 500 ng aflatrem (treatment group) to the heads of ants treated with 10% acetone solutions (control group). We found 261 genes that were significantly differentially expressed (Supplementary File 1). Of these genes, 148 were upregulated and 113 were downregulated in the treatment group (Figure 3). We found the largest increase of expression (∼22-fold increase) in a gene encoding an aminopeptidase N-like protein, a common cell surface hydrolase involved in a variety of cellular processes (Nocek et al., 2008). The largest decrease in expression (∼512-fold decrease) was exhibited by a gene coding for a cytochrome P450 6k1, a protein involved in detoxification of insect cells (Xing et al., 2021) (Figure 3). A GO enrichment analysis of these 261 genes identified three significantly over expressed GO terms, all three of which are broad categories: “sequence-specific DNA binding” (GO:0043565, adj. p-value 0.006), “DNA-binding transcription factor activity” (GO:0003700, adj. p-value 0.006), and “regulation of transcription, DNA-templated” (GO:00635, adj. p-value 0.048).

**Figure 3.**
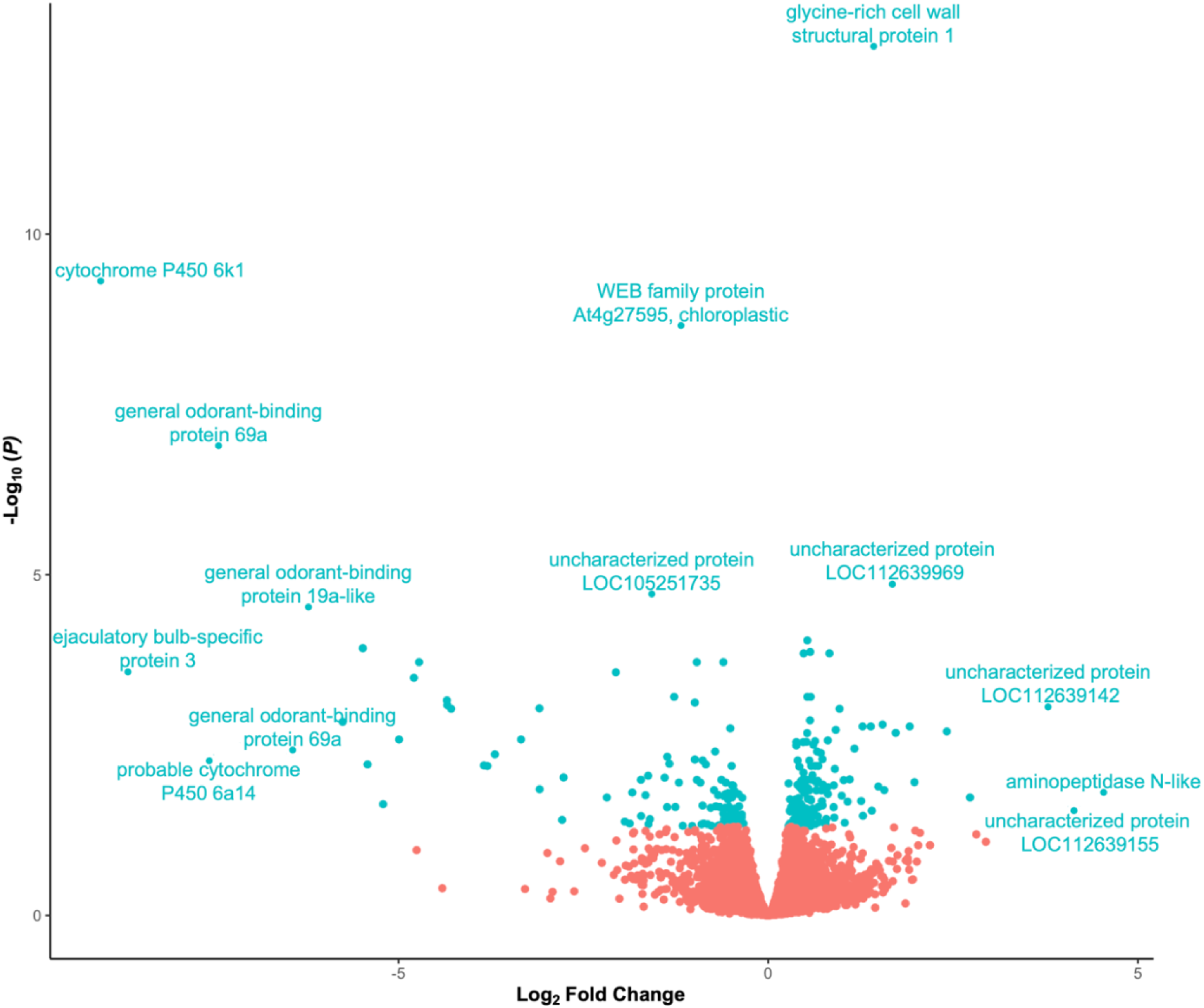
Volcano plot for the differential expression of 10,443 *Camponotus floridanus* genes in ants treated with 500 ng aflatrem compared to the 10% acetone control group. Dots in red represent non-significant changes in gene expression, while dots in blue represent significant changes using an alpha value of 0.05. Functional annotations are shown for genes with adjusted *P* values less than 5e-5 and absolute log_2_ fold changes lower than -6 or greater than 3.5.

Several of the 261 differentially expressed genes were predicted to play a role in neuromuscular systems. Among them, we found an *anoctamin-4*, which codes for an ion channel protein involved in Ca^2+^-dependent conductance (Reichhart et al., 2019), downregulated in aflatrem-treated ants. We also detected a *membrane metallo-endopeptidase* (*Neprilysin*), a transmembrane protein found in neurological tissue that plays a role in tissue perception and preservation (Obulesu, 2019), which was slightly upregulated in aflatrem-treated ants. More specific to muscular function, we found 2 upregulated *titins* genes, which code for very large proteins that play an essential role in sarcomere function (Freiburg et al., 2000; Zhang et al., 2000). Also upregulated was a *ryanodine receptor* gene, responsible for releasing Ca^2+^ in the smooth endoplasmic reticulum of muscle cells during excitation-contraction coupling (Lanner et al., 2010; Shakiryanova et al., 2007). On the other hand, *caveolin-3*, which codes for a scaffold protein that organizes signalling molecules in the membranes surrounding muscle cells (Dewulf et al., 2019; Galbiati & Lisanti, 2000-2013) was found to be downregulated, as well as 2 *myosin ID heavy chain-like protein* genes that act as the motor proteins of muscles (Wells et al., 1996) (Supplementary File 1).

In addition to neuromuscular genes, we also identified many downregulated olfactory-related genes, including 4 general odorant-binding protein genes (OBPs) (i.e., *OBP19a*, *OBP56a*, and 2 *OBP69a*). These proteins are located in the antennal sensillary lymph of insects and bind to pheromones that play a role in insect communication and interactions (Gaubert et al., 2020). Moreover, we found 3 significantly downregulated *ejaculatory bulb-specific* genes, involved in the release of pheromones in insects (Pelosi et al., 2018), and a downregulated *sensory neuron membrane protein* gene, which has been found to play a role in pheromone sensitivity in the moth *Heliothis virescens* (Pregitzer et al., 2014).

We also detected a differentially expressed gene coding for a circadian clock-controlled protein. Such genes could be of interest, given previous explorations of the time-dependent nature of behavioural manipulation and the disruption of clock-dependent foraging behaviours in *Ophiocordyceps*-infected ants (de Bekker et al., 2015; Das & de Bekker, 2022; Will et al., 2020), This gene, which showed an 18-fold decrease in expression in aflatrem-injected ants compared to 10% acetone controls, also contains a Haemolymph Juvenile Hormone Binding Protein Pfam domain. Such binding proteins, driven by circadian rhythms, are known to act as carrier proteins, chaperoning juvenile hormones, which play a wide array of roles in insect physiology and development, to their target tissues (Zalewska et. Al., 2009).

### 3.3 Comparison of aflatrem-induced DEGs with *Ophiocordyceps* infection DEGs

To investigate if any of the differentially expressed genes were affected during *Ophiocordyceps*-induced behavioural manipulation, we compared our RNA-seq results from the heads of aflatrem-injected ants to transcriptomics data obtained from the heads of *C. floridanus* manipulated by *Ophiocordyceps* (Will et al., 2020). We found that 113 of the 261 differentially expressed genes observed in aflatrem-treated ants were also differentially regulated in late-stage *Ophiocordyceps* infections (Supplementary File 1). Of these genes, 30 were differentially regulated in the same direction, while 83 shared an inverse relationship (Figure 4).

**Figure 4.**
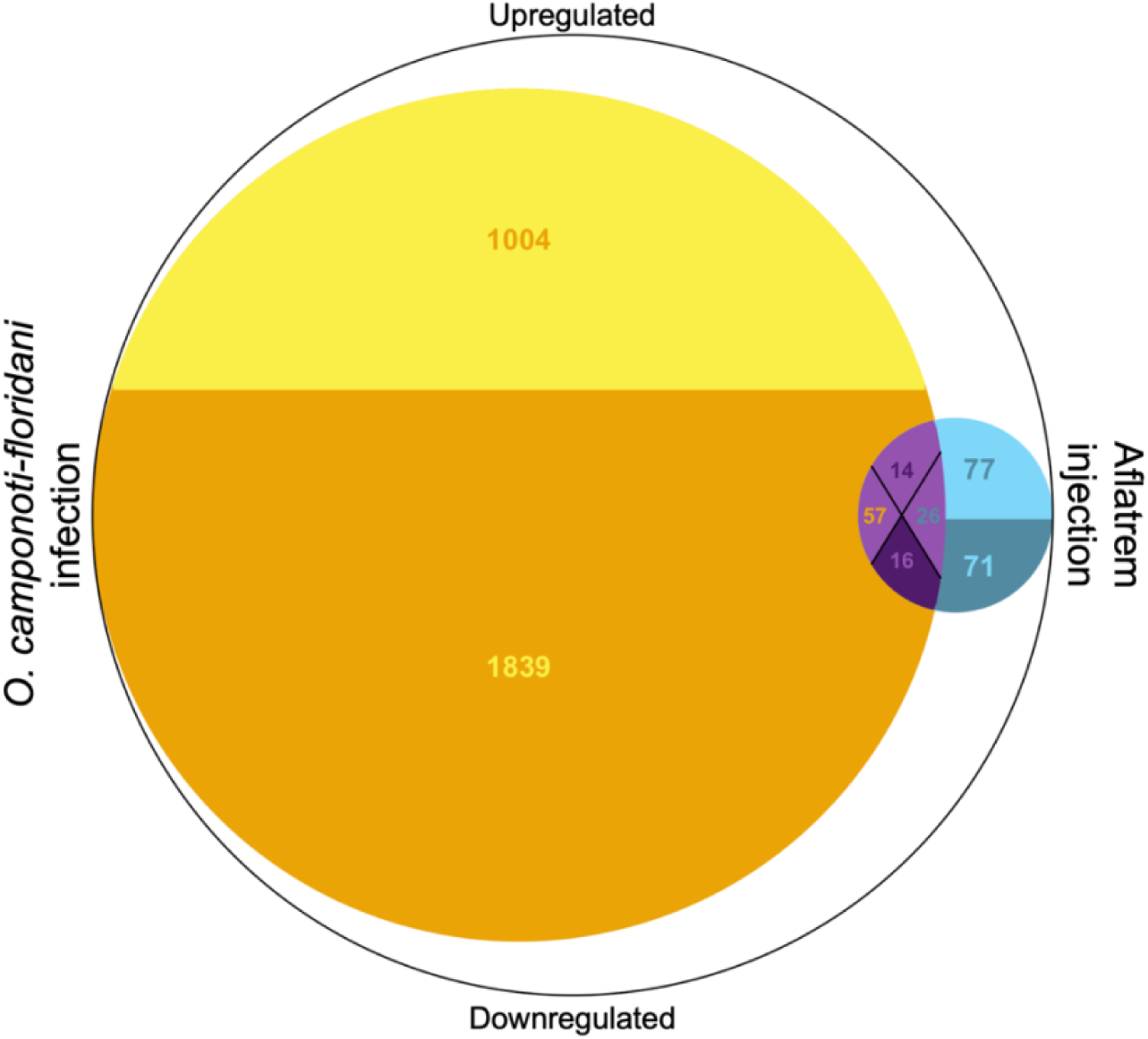
A modified Venn Diagram (i.e., a “Kaleidoscope Diagram”) representing the overlap and differences between the differentially expressed genes (DEGs) in the heads of *Ophiocordyceps-*infected ants (left) (Will et al., 2020) and the DEGs in the heads of aflatrem-treated ants (right) (this study), as compared to gene expression in their respective control groups. The lighter top colours represent upregulated genes while the darker bottom colours represent downregulated genes. In the centre overlapping region, the lighter north quadrant and darker south quadrant represent DEGs that are dysregulated in the same direction in both groups, either up or down respectively. Conversely, the west quadrant with orange texts represents DEGs with opposite expression from the perspective of upregulated *Ophiocordyceps-*infection genes and downregulated aflatrem-injection genes. The east quadrant with blue texts represents DEGs with opposite expression from the perspective of upregulated aflatrem-injection genes and downregulated *Ophiocordyceps-*infection genes.

The previously mentioned *cytochrome P450*, *sensory neuron membrane protein* encoding gene, *OBP56a*, and *caveolin-3* genes were amongst the similarly expressed genes, all of which were downregulated. We also found an upregulated *nuclear hormone receptor E75*, member of a family of transcription factors that are known to be activated by juvenile hormone signalling pathways in Diptera and Lepidoptera (Dubrovskaya et al., 2004). In addition, we detected an upregulated circadian clock steroid-producing gene linked to the maintenance of circadian rhythms during stress in *Drosophila* (Kumar S et al., 2014). Moreover, a *neurogenic locus protein delta*, capable of coding for multiple translational products including neural differentiation factors and growth hormones (Kopczynski et al., 1988), was also upregulated in both datasets.

Amongst the differently expressed genes, the aforementioned *OBP19a-like* gene, 5 *cytochrome P450s*, and an *ejaculatory bulb-specific protein 3*, were all significantly downregulated in aflatrem-treated ants but upregulated in *Ophiocordyceps*-infected ants. More genes shared this same inverse relationship, including an *odorant receptor coreceptor* gene, coding for proteins expressed in chemosensory organs that form heteromultimeric complexes within odorant receptors (Mukunda et al., 2014). We also detected a *short neuropeptide F*, which is implicated in the regulation of animal behaviour including circadian rhythms and hunger (Lee et al., 2004), and a *netrin receptor UNC5C*, which aids in the formation of axon connections through axon extension and cell migration (Kim & Ackerman, 2011). S*ialin*, a sialic acid transporter-coding gene that prevents neurodegenerative disorders caused by accumulation of sialic acid (Tarailo-Graovac et al., 2017) was additionally among these differentially expressed genes.

Upregulated in our aflatrem-treated ants, but downregulated in the *Ophiocordyceps*-infected ants, was a *receptor-type tyrosine-protein phosphatase delta* (*PTPRD*). This gene is part of the PTP family involved in many known signalling pathways for which delta is shown to promote neurite growth and regulate axon guidance (Tomita et al., 2020). In addition, we found an *obscurin*, which, like *titin*, codes for a large protein expressed in skeletal muscles that anchors myofibrils to the sarcoplasmic reticulum for muscle contraction (Feher, 2017), and a *juvenile hormone acid O-methyltransferase-like* gene, which codes for an enzyme that converts precursor molecules into juvenile hormones (Shinoda & Itoyama, 2003). Finally, the two data sets also shared 26 genes with statistically significant changes in expression coding for proteins of unknown function (Supplemental File 1).

## 4. DISCUSSION

Many parasites from across the Tree of Life utilize behavioural manipulation to increase their reproduction, survivability, and transmission at a cost to the host. Examples include *Loxothylacus panopaei* infecting crabs (Blakeslee et al., 2021), *Spinochordodes tellinii* nematomorphs infecting katydids (Biron et al., 2005), the protozoan *Toxoplasma gondii* infecting mice (Tong et al., 2021), *Cotesia congregata* wasps parasitizing caterpillars (Adamo, 2019), the virus *Rabies lyssavirus* infecting vertebrate animals (Hueffer et al., 2017), and *Entomophthora muscae* fungi infecting flies (Elya and Licht, 2021). Despite the growing awareness of this phenomena, the fundamental processes in which such behavioural manipulations are established, both in terms of the parasite’s role and the pathways affected within their animal hosts, remain poorly understood. In recent years, multi-omics work on “zombie ants” has provided a myriad of candidate compounds that *Ophiocordyceps* fungi might use to manipulate ant behaviour. However, their causal relationships with manipulated host behaviours still need to be functionally tested. This study aimed to test aflatrem-like compounds previously implicated in the establishment of *Ophiocordyceps* extended phenotypes in zombie ants and known to cause neuromuscular impairment in vertebrates (Selala et al., 2008; Valdes et al., 1985; Will et al., 2020). This “drunkard’s walk” behaviour has not been described in ants infected by other disease-causing agents (e.g., *Beauveria bassiana*) (Trinh et al., 2021), making it less likely that staggering should be considered a general sickness behaviour. Understanding the transferability of behaviour-affecting compounds and their effects at the genetic level is a step forward in elucidating the mechanisms underlying behavioural manipulation phenotypes and for determining whether neurological pathways affected during infection are conserved in nature. Moreover, learning how parasites can dysregulate host behaviour will provide insights into the regulation of animal behaviour in general.

We tested both the behavioural and genetic effects of aflatrem on carpenter ants by exposing healthy *C. floridanus* ants to high purity aflatrem and quantifying its behavioural effects. Following injections, we found dose of aflatrem to be positively correlated to the amount of time that treated individuals spent staggering. To rule out that this observed behaviour was a symptom related to death, we performed a longevity assay to demonstrate that aflatrem does not kill *C. floridanus*. These preliminary tests demonstrated that aflatrem-like compounds are unlikely to be employed by *Ophiocordyceps* species as a means to kill their hosts. Instead, the observed staggering behaviours indicate that the aflatrem-like compounds produced by *Ophiocordyceps* during infection are, at least in part, responsible for the drunkard’s walk phenotype exhibited by infected carpenter ants (Hughes et al., 2011; Trinh et al., 2021).

In vertebrates, staggers syndrome results from the potentiation of GABA chloride currents (Yao et al., 1989). Given that these receptors are conserved in ant species (Wnuk et al., 2014), it is plausible that similar mechanisms are at play here. Additionally, our RNA-Seq results suggest dysregulation of several genes that could explain dysfunction in neuromuscular activity. Among them, we found *anoctamin-4*, which was downregulated in aflatrem-treated ants and encodes a Ca^2+^-dependent non-selective monovalent cation channel (Reichhart et al., 2019). These channels play a vital role in regulating cation currents that drive excitation of neuronal and muscular activity (Hartzell et al. 2004). Likewise, *ryanodine receptor* also plays a role in cation regulation by releasing Ca^2+^ ions during contraction in skeletal muscles (Lanner et al., 2010). Dysregulation in both genes could impact proper muscle contraction in aflatrem-treated ants, resulting in difficulty walking. Furthermore, *caveolin-3*, a gene coding for structural components of the plasma membrane that regulate signal transduction events in skeletal muscle cells (Galbiati & Lisanti, 2000-2013), was also dysregulated. Mutations in caveolin-3 proteins are known to cause a variety of different muscle diseases in humans (Galbiati & Lisanti, 2000-2013), including caveolinopathy and distal myopathy. Both diseases are characterized by the onset of muscle weakness (Aboumousa et al., 2008). Moreover, caveolin-3-related rippling muscle disease is a disorder that can lead to repeated rapid contraction in muscles, particularly proximal muscles (Torbergsen, 2002). As such, *caveolin-3* dysregulation in aflatrem-treated ants could have given rise to the staggers that we observed. Our findings provide additional insight into the downstream effects on gene expression resulting from GABA potentiation as our RNA-Seq data reveals genes that may be associated with neuromuscular dysfunction as a result of aflatrem-induced staggering. Given the similarity of effects observed between our finding with invertebrates and previous work in cows, our data could be informative for future research in vertebrate models.

Our experiments further demonstrated that aflatrem has a significant effect on the mobility of ants, decreasing overall activity as dose increased. Ants injected with higher doses of aflatrem moved less often, did so much slower when they were active, and as a result travelled shorter distances. We also observed that they had more difficulties climbing, particularly when it involved the twine in the arena which requires greater dexterity than climbing on the wall or top of the petri dish. Previous field studies suggest that *Ophiocordyceps* development and fruiting body formation, needed for spore transmission, are dependent on precise levels of light, humidity and temperature (Andersen & Hughes, 2012; Andriolli et al., 2019; Cardoso Neto et al., 2019; Hughes et al., 2011; Lavery et al., 2021; Will et al., under review). The reduced speed and disruption of normal gate induced by the upregulation of aflatrem-like compounds during late infection could aid to keep the ant within these optimal microclimates prior to biting, by preventing it from climbing too high into the canopy. However, further studies would be needed to test this hypothesis as lethargy is a universally common sickness behaviour as well.

While we could reason a parasite-adaptive function for aflatrem-reduced activity, it seemingly contradicts previous studies where *Ophiocordyceps*-infected individuals showed hyperactive locomotion prior to summiting (de Bekker et al., 2015; Trinh et al., 2021; Will et al., 2020). Nevertheless, transcriptomics studies on *Ophiocordyceps*-manipulated ants suggest that the fungus expresses manipulation compounds in a time-specific manner (de Bekker et al., 2015; Will et al., 2020). It is reasonable to suggest that, prior to the upregulation of aflatrem-like compounds, *Ophiocordyceps* secretes effectors that induce hyperactivity and climbing behaviours to drive the ant away from the nest and up nearby structures. An example of one such effector could be protein tyrosine phosphatase (PTP). Genes encoding this protein are upregulated during *Ophiocordyceps* manipulation (de Bekker et al., 2015; Will et al., 2020) and have been found to induce hyperactivity and enhanced locomotion in caterpillars that are infected with a behaviour-manipulating baculovirus (Han et al., 2015). Enhanced locomotion could be parasite-adaptive by leading hosts away from aggressive conspecifics, which can recognize infected individuals and attack them (Trinh et al., 2021; Will et al., 2020) as part of their social immunity (Cremer et al., 2007). After the ant is safely away from the adaptive behaviours of the colony, the production of debilitating compounds could function to maintain the ant in a desirable, elevated location.

In addition, aflatrem-like compounds may also aid in the positioning of the host for the final, and perhaps most iconic, behavioural manipulation; the “death grip”. During the death grip, ants bite onto vegetation with their mandibles, firmly anchoring them in place. Shortly after, the fungus kills the host and quickly consumes the remaining nutrients to produce its reproductive structures. The attachment of the host is a vital step in host manipulation as it promotes infective spore formation and release. Without it, ants would fall back to the forest floor, which interferes with fruiting body formation and, eventually, transmission (Andersen et al., 2012; Andriolli et al., 2019; Loreto et al., 2014; Will et al., under revision). The large increase in resting behaviours seen in ants that received the highest dose of aflatrem could aid in positioning of the ant’s head for biting behaviours. Moreover, aflatrem injection resulted in the differential expression of muscle-related genes in the heads of treated ants as compared to the control group. Therefore, the muscle-related effects of aflatrem-like compounds secreted by *Ophiocordyceps* could potentially be directly involved in the locking of the mandible muscles as part of the biting phenotype (Hughes et al., 2011). However, we did not observe increased biting behaviour in aflatrem-treated ants in this study. This indicates that if aflatrem-like compounds do play a role in maintaining optimal positioning for adherence, it requires the aid of other compounds to induce the death grip phenotype.

As part of the death grip phenotype, *Ophiocordyceps*-infected *C. floridanus* are often found wrapping their limbs around thin pieces of Spanish moss, which they most frequently use as a biting substrate (Figure 5) (Will et al., under review). This “hugging” behaviour has also been observed in other *Ophiocordyceps*-infected ants where smooth substrates, such as twigs, are primarily bitten (Loreto et al., 2018). The transcription-level dysregulation of sensory perception and neuromuscular functioning that we found, as well as the observed staggers, suggest that unsteadiness during climbing might cause ants to stumble and hug the vegetation. As such, the hugging behaviour itself could act as a secondary means by which hosts are kept in elevated positions should biting itself fail. Indeed, the adherence to vegetation through biting is sometimes missing in *Ophiocordyceps*-manipulated ants, both in the field and in the lab (Will et al., 2020) while hugging is always observed. In fact, our group has witnessed multiple examples of such instances in the field where hosts with fully grown fruiting bodies are held to strands of Spanish moss by just their limbs, either due to failed biting or the complete loss of the ant’s head from the sample. Hugging may be unique to ant hosts of *Ophiocordyceps* species as the phenotype is absent from other host-manipulating entomopathogens, such as *Eryniopsis lampyridarum* infecting the Goldenrod Soldier Beetle, which similarly induces summiting and death grip behaviours on thin, huggable foliage (Steinkraus et al., 2017).

**Figure 5.**
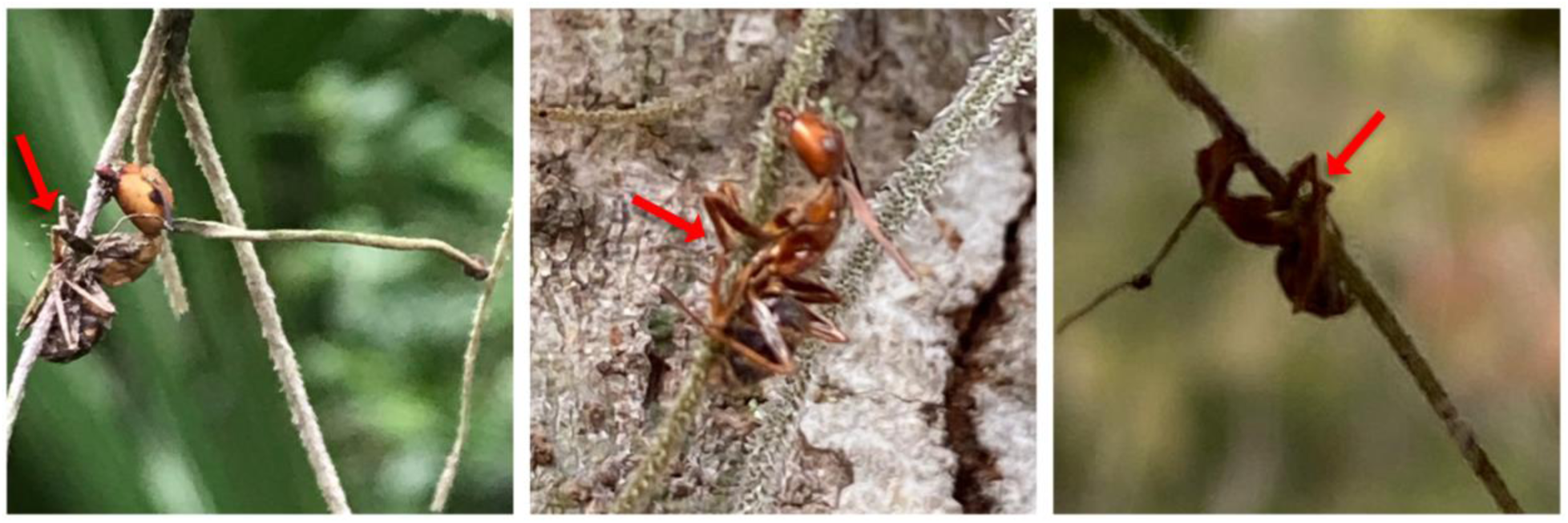
Examples of “hugging” behaviour portrayed by *Ophiocordyceps camponoti-floridani*-infected *Camponotus floridanus* ants during the death grip observed in a natural setting.

While previous genomic analyses have demonstrated that the secondary metabolite cluster responsible for the production of aflatrem-like compounds is conserved across *Ophiocordyceps* species infecting members of the Camponotini tribe, the exact structure and concentrations of these compounds produced *in vivo* are still unknown. Regardless, we were able to use reasonable doses of purified aflatrem (i.e., within body weight equivalents used in previous pesticide testing) to replicate the staggers/“drunkard’s walk” phenotype induced in ants during *Ophiocordyceps* infection. This caused the differential expression of 108 ant host genes that were previously identified in *Ophiocordyceps*-infected ants exhibiting the summiting phenotype. As such, at least in part, similar host pathways appear to be targeted during isolated aflatrem-dosing and *Ophiocordyceps* infection, leading to comparable behaviours. This indicates that aflatrem-like compounds secreted by *Ophiocordyceps* species are likely responsible for some of the transcriptional and behavioural changes observed during manipulation. Future work characterizing the chemical composition and three-dimensional structure of aflatrem-like compounds produced by species of the *O. unilateralis* complex is needed to shed further light on the physiological effects of these alkaloids and their role in behaviour manipulation. Nevertheless, our work demonstrates that exposing hosts to hypothesized parasite manipulation compounds is a valuable avenue to test their proposed effects at the genome and phenome level, and establishes a framework for future studies characterizing the effects of host-manipulating compounds. Furthermore, these sorts of in vitro studies provide a baseline for the comparison of host pathways affected by other manipulating parasites to determine if other systems share a common approach towards manipulation. As such, integrative behavioural studies like this one bring us one step closer to characterizing the molecular mechanisms that drive not only zombie-making fungi, but other behavior-manipulating parasites across the Tree of Life.

## AKNOWLEDGMENTS

We would like to extend our thanks to the King lab for their advice on using, and access to, their ShopVac as a method for collecting ants, Biplabendu Das for his recommendations and assistance with the CowLog3 and MATLAB programs, and Ian G. Will for his knowledgeable insight into the Ophiocordyceps ergot alkaloid gene cluster and advice on statistical analysis of our data. The concept for the Kaleidoscope diagram was designed by William C. Beckerson. We would like to thank Andrew Swafford for building an R-package that generated the Kaleidoscope diagram we used in this manuscript.

This work is supported by NSF PRFB Award 2109435 to William C. Beckerson and NSF CAREER Award 1941546 to Charissa de Bekker.

The raw RNA-Seq data used in these analyses can be found at NCBI’s Sequence Read Archive under BioProject ID PRJNA914723 https://www.ncbi.nlm.nih.gov/sra/PRJNA914723

The raw CowLog observation data can be accessed at: https://github.com/WCBeckerson/28-Minutes-Later-Data

## AUTHOR CONTRIBUTIONS

WCB and CdB designed the experiment, analysed the data, performed the RNA extractions, and wrote the paper. WCB, CK, and UAM collected the ants, performed the injections, and recorded the video footage. CK and UAM performed the behavioural analyses using CowLog. WCB ran the MARGO software and performed the statistical analyses.

## Supplemental Files

**Supplemental Table 1:** The average length and weight of *Camponotus floridanus* in this study

| Colony # | Treatment Group | Average Length (mm) | Average Weight (mg) |
| --- | --- | --- | --- |
| Colony 1 | C1-PBS | 7.07 | 7.56 |
|  | C1-PBS/10% acetone | 6.50 | 6.93 |
|  | C1-PBS/10% acetone/500ng aflatrem | 6.78 | 7.94 |
|  | C1-PBS/10% acetone/100ng aflatrem | 7.00 | 8.14 |
|  | C1-PBS/10% acetone/50ng aflatrem | 6.43 | 8.44 |
|  | C1-PBS/10% acetone/10ng aflatrem | 7.29 | 9.01 |
| Colony 2 | C2-PBS | 7.07 | 9.49 |
|  | C2-PBS/10% acetone | 7.07 | 10.27 |
|  | C2-PBS/10% acetone/500ng aflatrem | 6.93 | 11.09 |
|  | C2-PBS/10% acetone/100ng aflatrem | 6.93 | 10.48 |
|  | C2-PBS/10% acetone/50ng aflatrem | 6.86 | 10.86 |
|  | C2-PBS/10% acetone/10ng aflatrem | 6.86 | 10.13 |
| Colony 3 | C3-PBS | 6.07 | 5.40 |
|  | C3-PBS/10% acetone | 6.93 | 7.93 |
|  | C3-PBS/10% acetone/500ng aflatrem | 6.79 | 6.44 |
|  | C3-PBS/10% acetone/100ng aflatrem | 5.71 | 5.40 |
|  | C3-PBS/10% acetone/50ng aflatrem | 5.64 | 6.23 |
|  | C3-PBS/10% acetone/10ng aflatrem | 6.29 | 7.24 |
|  | Total | 6.68 mm | 8.28 mg |

**Supplemental Figure 1.**
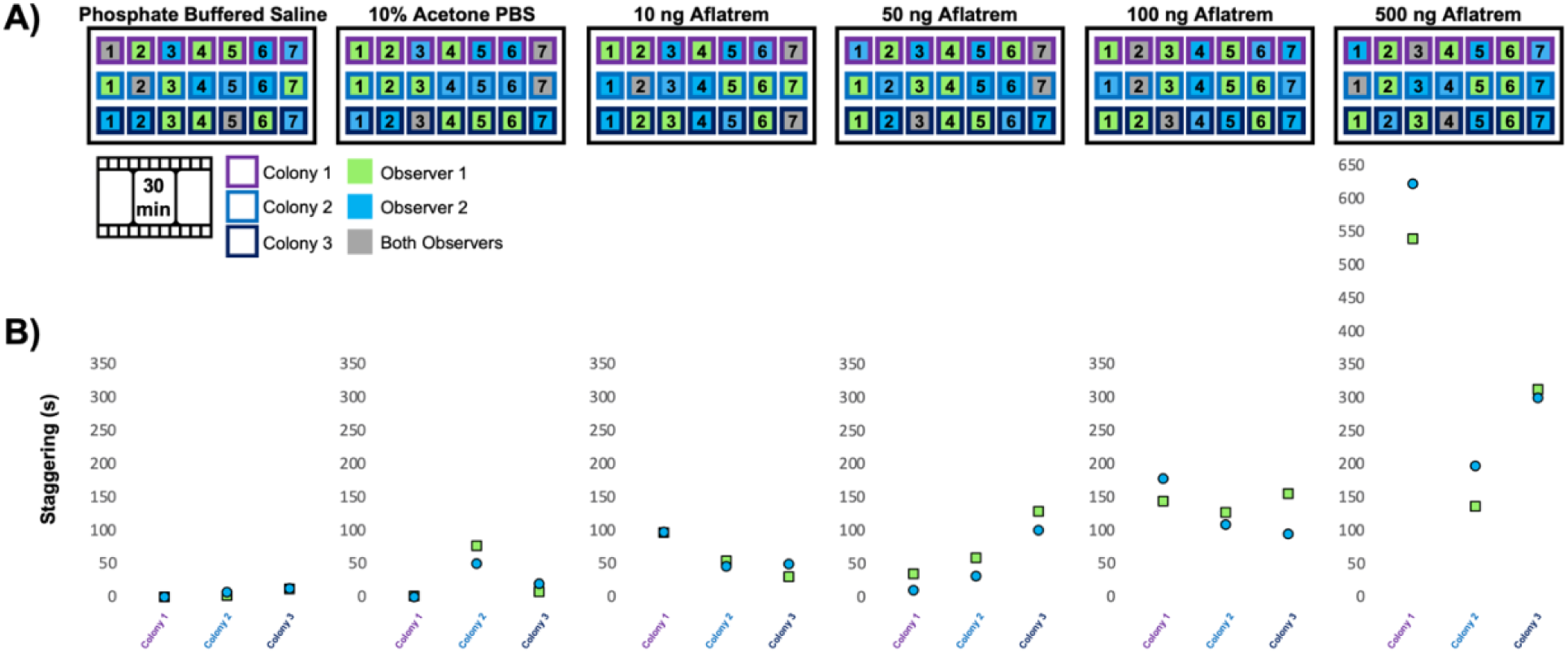
An overview of the randomized block format used to assign videos to each observer (A). The videos assigned to observer 1 are shown in green, the videos assigned to observer 2 are shown in blue, and the videos assigned to both for formative analysis are shown in grey. B) depicts the duration of staggering scores for each grey assignment, with the scores for observer 1 shown as green squares and the scores for observer 2 shown as blue circles.

**Supplemental Figure 2.**
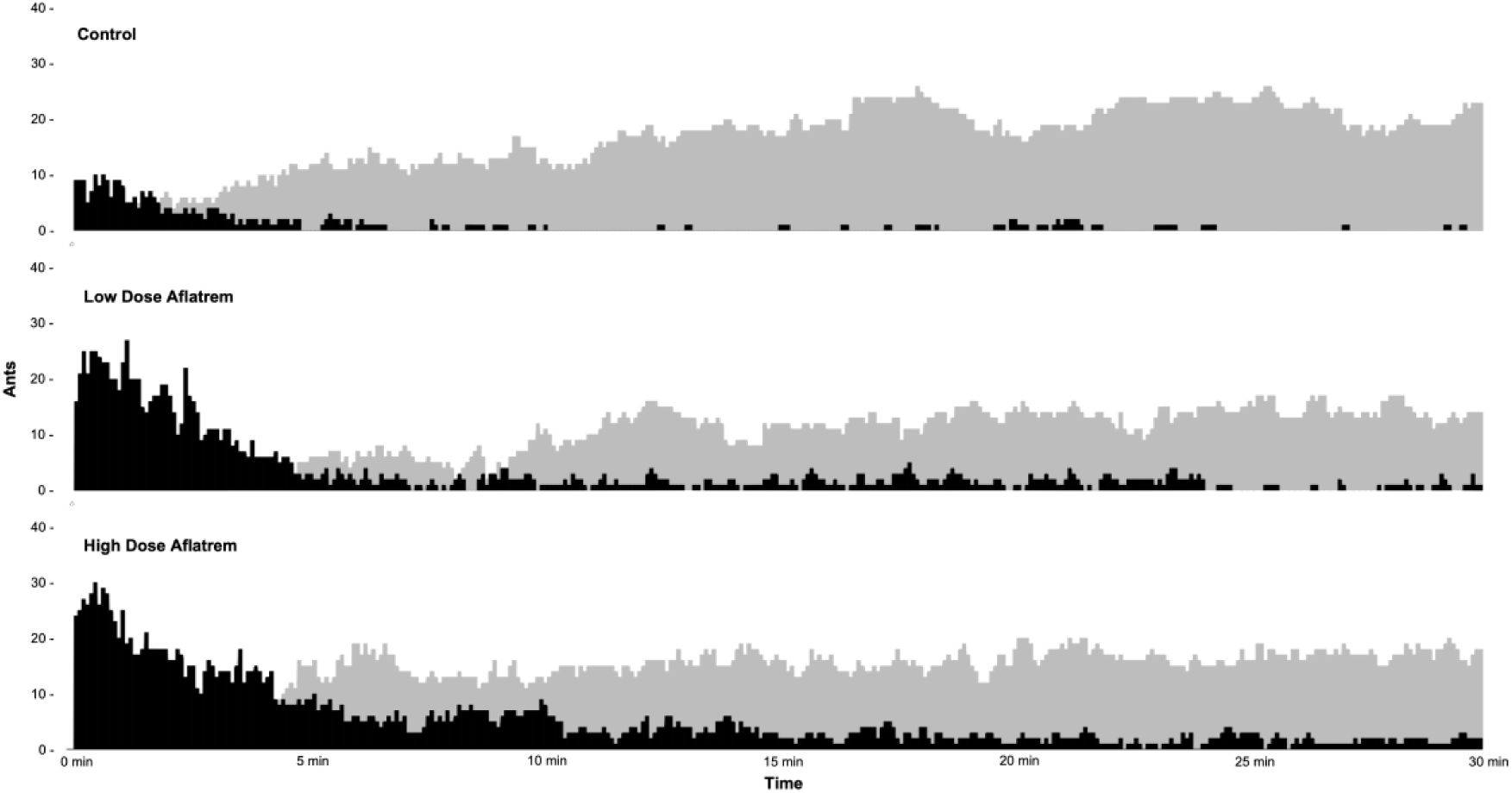
Bar graphs depict the number of ants exhibiting staggering behaviour (black) and the number of ants resting (dark grey) across the 30-minute observational period, broken down into 5 second intervals. Control refers to ants injected with 10% acetone solvent. Low dose aflatrem shows the data for ants that received 10-50 ng aflatrem, while high dose aflatrem visualizes staggering and resting for ants that received 100-500 ng.

**Supplemental Figure 3.**
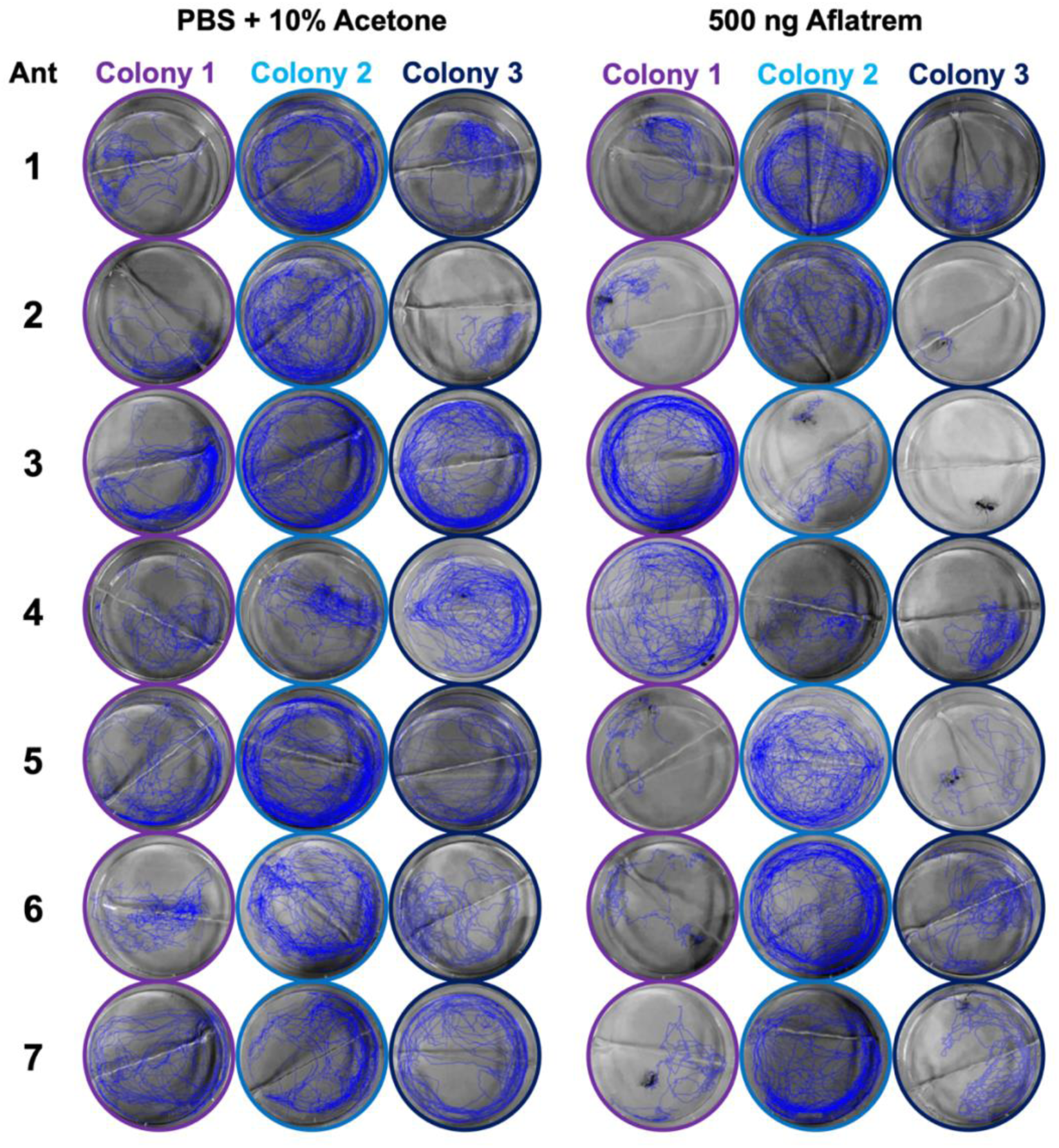
The path travelled by each ant recorded by MARGO tracking software during the 30-minute observational period is shown in dark blue. The three columns on the left show the paths travelled by the control group injected with 10% acetone, while the three columns on the right show the paths travelled by ants in the 500 ng aflatrem treatment group. Each group is further divided by colony ID using purple, blue, and dark blue outline to indicate individuals from colonies 1, 2, and 3, respectively.

**Supplemental Figure 4.**
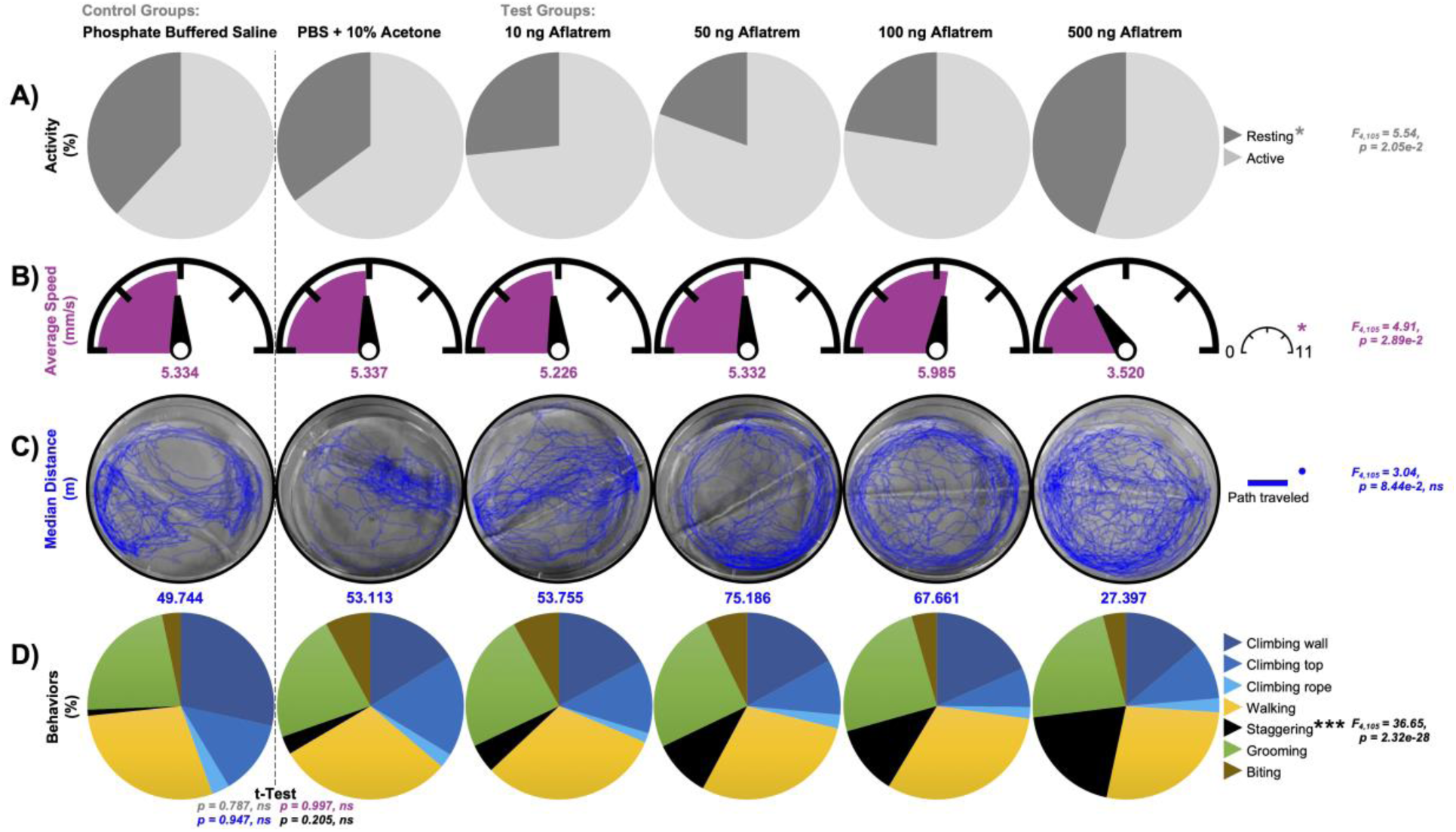
A graphical overview of the relative time spent resting (A), average speed calculated by MARGO (B), the path travelled for the median distance in each group (C), and the distribution of active time spent in the 7 commonly observed behaviours (D). Column 1 represents a pure Phosphate Buffered Saline (PBS) control, column 2 represents the 0 dose control containing 10% acetone without Aflatrem, and columns 3-6 represent the dose assay from 10 – 500 ng of Aflatrem.

Supplemental File 1 *\*Raw reads from the RNA-Seq data and summary table of the 261 significantly dysregulated genes compared to results from Will, et al., 2020*

**Supplemental Video 1 – mp4.**
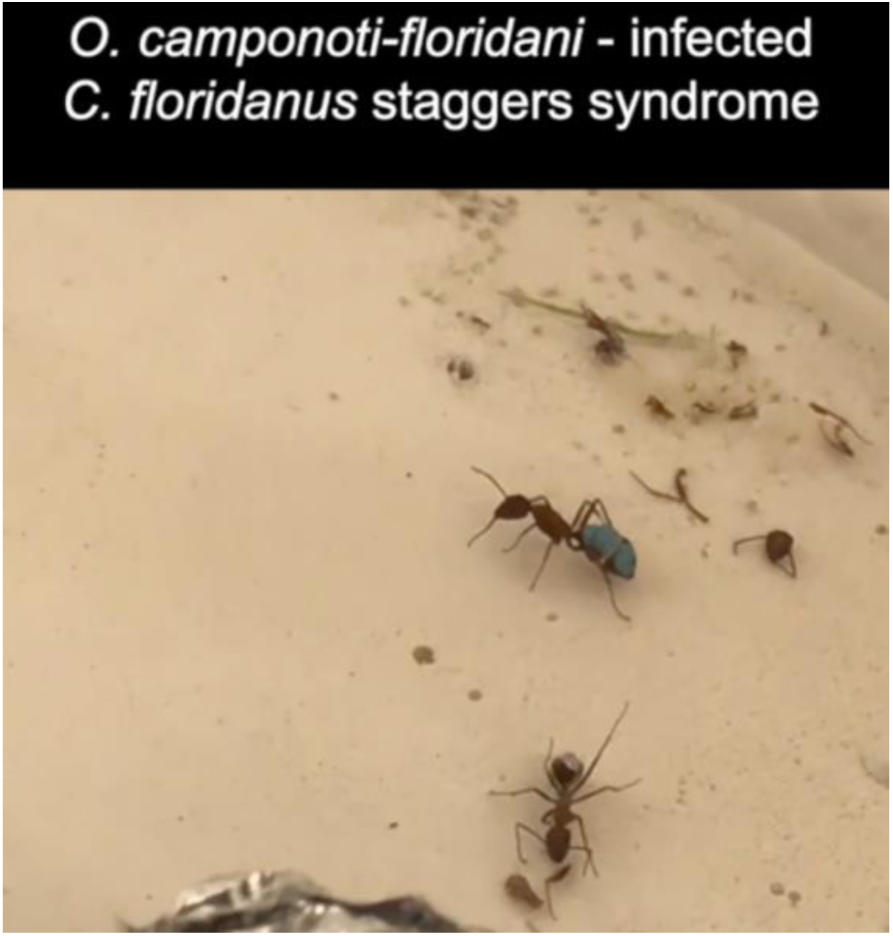
\**Ophiocordyceps camponoti-floridani* – infected *Camponotus floridanus* exhibiting the “drunkard’s walk” reminiscent of staggers syndrome observed in vertebrates exposed to aflatrem.

**Supplemental Video 2 – mp4.**
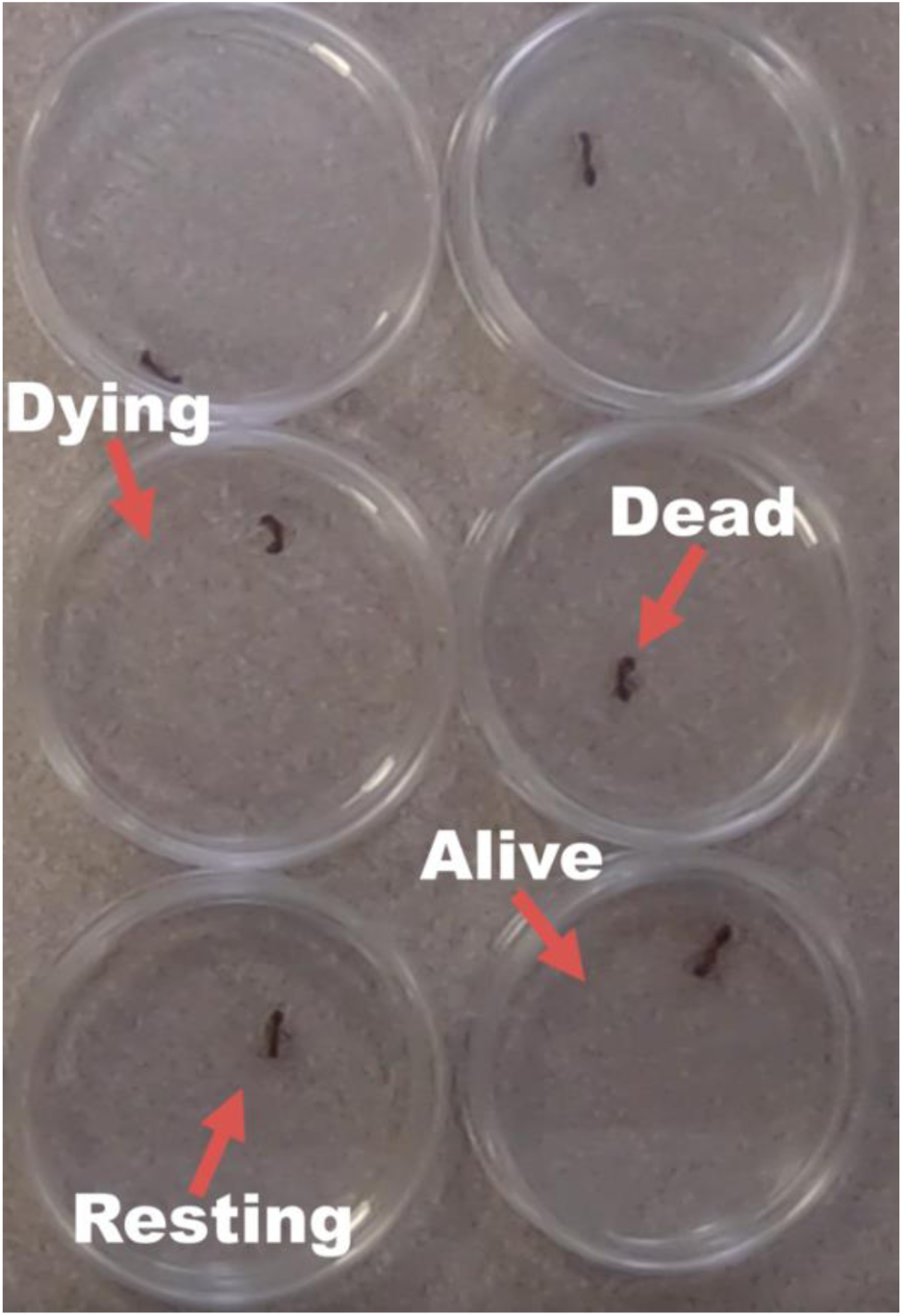
*A timelapse video of the preliminary test for acetone tolerance. This clip demonstrates the body positions of ants that are alive, resting, dying, and dead.

**Supplemental Video 3 – mp4.**
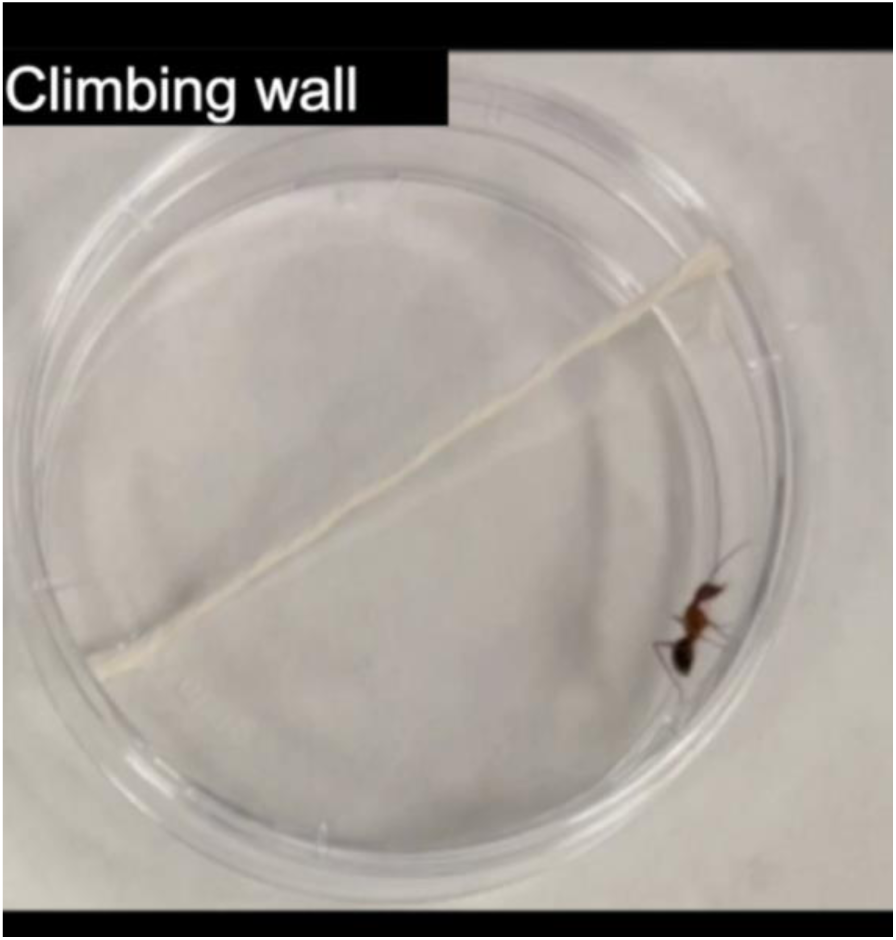
*The eight clearly defined behavioural phenotypes observed and scored using CowLog3. These included climbing the wall, climbing on the top, climbing the rope, walking, staggering, grooming, resting, and biting.

**Supplemental Video 4 – mp4.**
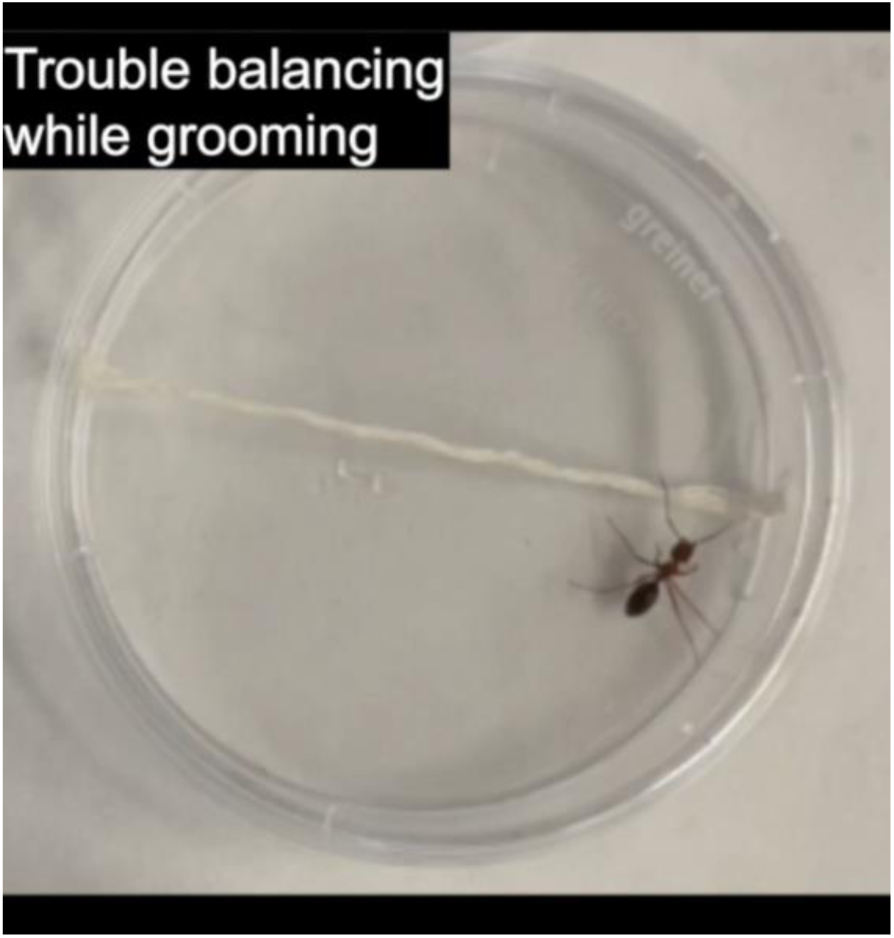
* The six subcategories of staggering behaviours observed upon closer observation. These included trouble balancing while grooming, listing to one side, falling over, walking in tight circles, flipping over, and gaster wagging.

**Supplemental Video 5 – mp4.**
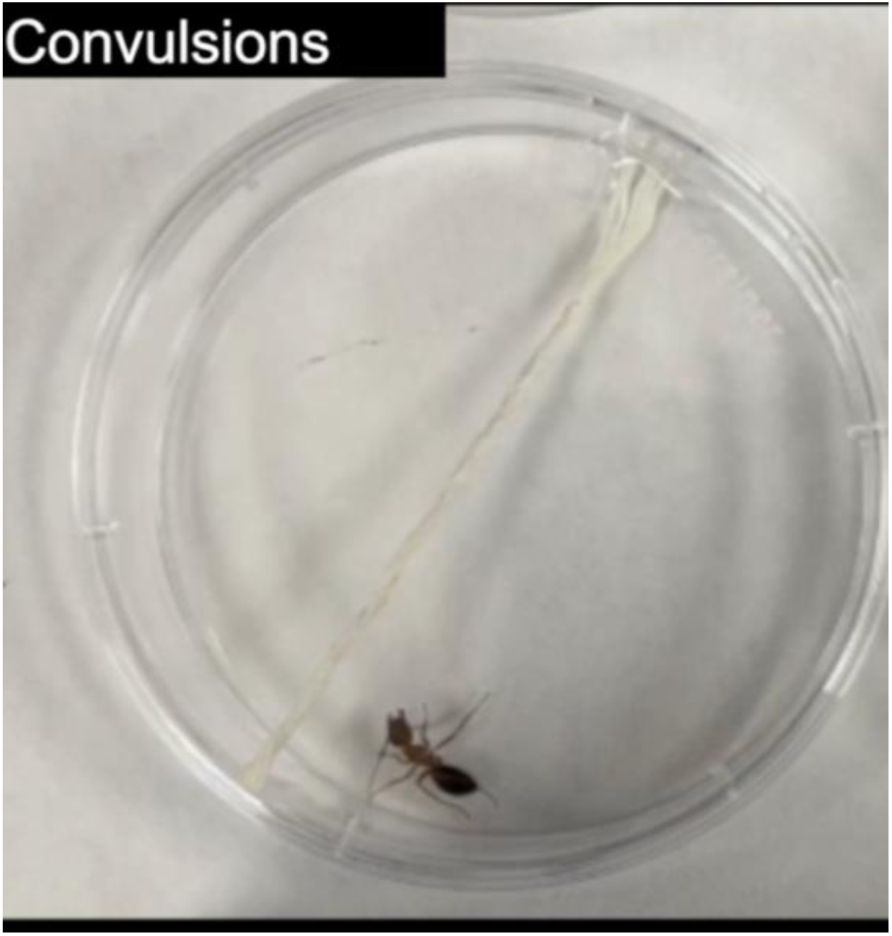
*Three miscellaneous behaviours observed in rare quantities during aflatrem exposure. These included walking backwards, lunging/bucking, and convulsions.

Supplemental R-package 1 *\*The code required to produce a Kaleidoscope diagram*

## REFERENCES

Aboumousa A, Hoogendijk J, Charlton R, Barresi R, Herrmann R, Voit T, Hudson J, Roberts M, Hilton-Jones D, Eagle M, Bushby K, Straub V. (2008). Caveolinopathy—new mutations and additional symptoms. Neuromuscular Disorders 18(7): 572–578. https://doi.org/10.1016/j.nmd.2008.05.003

Adamo SA. (2019). Turning your victim into a collaborator: Exploitation of insect behavioural control systems by parasitic manipulators. Current Opinion in Insect Science. 33: 25–29. https://doi.org/10.1016/j.cois.2019.01.004

Andersen SB, Hughes DA. (2012). Host specificity of parasite manipulation. Communicative & Integrative Biology 5(2), 163–165. https://doi.org/10.4161/cib.18712

Andersen SB, Gerritsma S, Yusah KM, Mayntz D, Hywel-Jones NL, Billen J, Boomsma JJ, Hughes DP. (2009). The life of a dead ant: the expression of an adaptive extended phenotype. The American Naturalist 174(3): 424–33. https://doi.org/10.1086/603640

Andrews S. (2010). FastQC: A Quality Control Tool for High Throughput Sequence Data [Online]. Available online at: http://www.bioinformatics.babraham.ac.uk/projects/fastqc/

Andriolli FS, Ishikawa NK, Vargas-Isla R, Cabral TS, de Bekker C, Baccaro FB. (2019). Do zombie ant fungi turn their hosts into light seekers? Behavioural Ecology 30(3), 609–616. https://doi.org/10.1093/beheco/ary198

Araújo JPM, Evans HC, Fernandes IO, Ishler MJ, Hughes DP. (2020). Zombie-ant fungi cross continents: II. *Myrmecophilous hymenostilboid* species and a novel zombie lineage. Mycologia 112(6): 1138–1170. https://doi.org/10.1080/00275514.2020.1822093

Araújo JPM, Hughes DP. (2019). Zombie-ant fungi emerged from non-manipulating, beetle-infecting ancestors. Current Biology 29(21): 3735–3738.e2. https://doi.org/10.1016/j.cub.2019.09.004

Araújo JPM, Evans HC, Kepler R, Hughes DP. (2018). Zombie-ant fungi across continents: 15 new species and new combinations within *Ophiocordyceps*. L. Myrmecophilous hirsutelloids species. Studies in Mycology 90: 119–160 https://doi.org/10.1016/j.simyco.2017.12.002

Arruda IDP, Villanueva-Bonilla GA, Faustino ML, Moura-Sobczak JCMS, Sobczak JF. (2021). Behavioural manipulation of the spider *Macrophyes pacoti* (Araneae: Anyphaenidae) by the araneopathogenic fungus *Gibellula* sp. (Hypocreales: Cordycipitaceae). Candian Journal of Zoology 99(5): 401–408. https://doi.org/10.1139/cjz-2020-0232

Biron DG, Marché L, Ponton F, Loxdale HD, Galéotti N, Renault L, Joly C, Thomas F. (2005). Behavioural manipulation in a grasshopper harbouring hairworm: a proteomics approach. Proceedings of the Royal Society of Biological Sciences 272(1577): 2117–2126. https://doi.org/10.1098/rspb.2005.3213

Blakeslee AMH, Pochtar DL, Fowler AE, Moore CS, Lee TS, Barnard RB, Swanson KM, Lukas LC, Ruocchio M, Torchin ME, Miller AW, Ruiz GM, Tepolt CK. (2021). Invasion of the body snatchers: the role of parasite introduction in host distribution and response to salinity in invaded estuaries. Proceedings of the Royal Society of Biological Sciences 288(1953): 288: 20210703. https://doi.org/10.1098/rspb.2021.0703

Bolger AM, Lohse M, Usadel B. (2014). Trimmomatic: A flexible trimmer for Illumina Sequence Data. *Bioinformatics*, btu170. https://doi.org/10.1093/bioinformatics/btu170

Breed MD, Moore J. (2010). The Encyclopedia of Animal Behavior. Academic Press.

Buczkowski G, Wossler TC. (2019). Controlling invasive Argentine ants, *Linepithema humile*, in conservation areas using horizontal insecticide transfer. Scientific Reports 9: 19495. https://doi.org/10.1038/s41598-019-56189-1

Cardoso Neto JA, Leal LC, Baccaro FB. (2019). Temporal and spatial gradients of humidity shape the occurrence and the behavioural manipulation of ants infected by entomopathogenic fungi in Central Amazon. Fungal Ecology 42, 100871. https://doi.org/10.1016/J.FUNECO.2019.100871

Chhetri P.K., Pradhan BK, Chhetri B. (2020) First record of *Ophiocordyceps dipterigena* Berk. & Broome (Ophiocordycipitaceae) in the Himalayas. National Academy Science Letters 43, 367–369. https://link.springer.com/article/10.1007/s40009-019-00857-3

Conesa A, Gotz S, Garcia-Gomez JM, Terol J, Talon M, Robles M. (2005) Blast2GO: a universal tool for annotation, visualization and analysis in functional genomics research. Bioinformatics 21: 3674–3676. https://doi.org/10.1093/bioinformatics/bti610

Cooley JR, Marshall DC, Hill KBR. 2018. A specialized fungal parasite (Massospora cicadina) hijacks the sexual signals of periodical cicadas (Hemiptera: Cicadidae: Magicicada). Scientific Reports 8:1432. https://doi.org/10.1038/s41598-018-19813-0

Cremer S, Armitage SAO, Schmid-Hempel P. (2007). Social Immunity. Current Biology 17(16) R693–R702. https://doi.org/10.1016/j.cub.2007.06.008

Danecek P, Bonfield JK, Liddle J, Marshall J, Ohan V, Pollard MO, Whitwham A, Keane T, McCarthy SA, Davies RM, Li H. (2021). Twelve years of SAMtools and BCFtools. GigaScience 10(2): giab008. https://doi.org/10.1093/gigascience/giab008

Das, B. (2022). timecourseRnaseq: an R package to analyze time-course RNAseq data. https://github.com/biplabendu/timecourseRnaseq

Das B, de Bekker C. (2022). Time-course RNASeq of *Camponotus floridanus* forager and nurse ant brains indicate links between plasticity in the biological clock and behavioural division of labor. BMC Genomics 23(57). https://doi.org/10.1186/s12864-021-08282-x

Dawkins R. (1982). The Extended Phenotype: The Long Reach of the Gene. Oxford, UK: Oxford University Press.

De Bekker C, Beckerson WC, Carolyn Elya. (2021). Mechanisms Behind the Madness: How do zombie-making fungal entomopathogens affect host behaviour to increase transmission? mBio 12(5), e01872–21.https://doi.org/10.1128/mBio.01872-21

de Bekker C, Ohm RA, Evans HC, Brachmann A, Hughes DP. (2017). Ant-infecting Ophiocordyceps genomes reveal a high diversity of potential behavioural manipulation genes and a possible major role for enterotoxins. Sci Rep 7:12508. https://doi.org/10.1038/s41598-017-12863-w

de Bekker C, Ohm RA, Loreto RG, Sebastian A, Albert I, Merrow M, Brachmann A, Hughes DP. (2015). Gene expression during zombie ant biting behaviour reflects the complexity underlying fungal parasitic behavioural manipulation. BMC Genomics 16:620. https://doi.org/10.1186/s12864-015-1812-x

Dewulf M, Köster DV, Sinha B, Viaris de Lesegno C, Chambon V, Bigot. A, Bensalah M, Negroni E, Tardif N, Podkalicka. J, Johannes L, Nassov P, Butler-Browne G, Lamaze C, Blouin CM. (2019). Dystrophy-associated caveolin-3 mutations reveal that caveolae couple IL6/STAT3 signaling with mechanosensing in human muscle cells. Nature Communications 10: 1974. https://doi.org/10.1038/s41467-019-09405-5

Dobin A, Davis CA, Schlesinger F, Drenkow J, Zaleski C, Jha S, Batut P, Chaisson M, Gingeras TR. (2013). STAR: ultrafast universal RNA-seq aligner. Bioinformatics. 29(1):15–21. https://doi.org/10.1093/bioinformatics/bts635

Dubrovskaya VA, Berger EM, Dubrovsky EB. (2004). Juvenile hormone regulation of the E75 nuclear receptor is conserved in Diptera and Lepidoptera. Gene 340(2): 171–177. https://doi.org/10.1016/j.gene.2004.07.022

Duran RM, Cary JW, Calvo AM. (2007). Production of cyclopiazonic acid, aflatrem, and aflatoxin by *Aspergillus flavus* is regulated by veA, a gene necessary for sclerotial formation. Applied Microbiology and Biotechnology 73(5): 1158–68. https://doi.org/10.1007/s00253-006-0581-5

Eldefrawi ME, Gant DB, Eldefrawi AT. (1990). The GABA receptor and the action of tremorgenic mycotoxins. Pohland AE (Ed.). Microbial Toxins in Foods and Feeds: Cellular and molecular modes of action (291-295). New York, NY, Plenum Press. https://doi.org/10.1007/978-1-4613-0663-4_27

Elya C, Lich HHdF. (2021). The genus Entomophthora: bringing the insect destroyers into the twenty-first century. IMA Fungus 12:34. https://doi.org/10.1186/s43008-021-00084-w

Evans HC, Araújo JPM, Halfeld VR, Hughes DP. Epitypification and re-description of the zombie-ant fungus, Ophiocordyces unilateralis (Ophiocordycipitaceae). Fungal Systematics and Evolution 1(1): 13–22. https://doi.org/10.3114/fuse.2018.01.02

Fain A. (1994) Adaptation, specificity and host-parasite coevolution in mites (Acari). International Journal for Parasitology 24(8): 1273–1283. https://doi.org/10.1016/0020-7519(94)90194-5

Feher J. (2017). Contractile Mechanisms in Skeletal Muscle. in: Quantitative Human Physiology pp. 305–217 :Academic Press. https://doi.org/10.1016/B978-0-12-800883-6.00028-8

Flor HH. (1956). The complementary genic systems in flax and flax rust. Advances in Genetics 8: 29–54. https://doi.org/10.1016/S0065-2660(08)60498-8

Fredericksen MA, Zhang Y, Hazen ML, Hughes DP. (2017). Three-dimensional visualization and a deep-learning model reveal complex fungal parasite networks in behaviourally manipulated ants. PNAS 114(*47*): 12590–12595. https://doi.org/10.1073/pnas.1711673114

Freiburg A, Trombitas K, Hell W, Cazorla O, Fougerousse F, Center T, Kolmerer B, Witt C, Beckmann JS, Gregorio CC, Granzier H, Labeit S. (2000). Series of exon-skipping events in the elastic spring region of titin as the structural basis for myofibrillar elastic diversity. Circulation Research 86(11): 1114–1121. https://doi.org/10.1161/01.res.86.11.1114

Galbiati F, Lisanti MP. (2000-2013). Caveolin-3 and Limb-Girdle Muscular Dystrophy. In: Madame Curie Bioscience Database [Internet]. Austin (TX): Landes Bioscience; 2000–2013. Available from: https://www.ncbi.nlm.nih.gov/books/NBK6073/

Gant DB, Cole RJ, Valdes JJ, Eldefrawi ME, and Eldefrawi AT. (1987). Action of tremorgenic mycotoxins on GABAA receptor. Life Sciences 41: 2207–2214. https://doi.org/10.1016/0024-3205(87)90517-0

Gaubert A, Amigues B, Spinelli S, Cambillau C. (2020). Chapter Seven – Strucutre of odorant binding proteins and chemosensory proteins determined by X-ray crystallography. Methods in Enzymology 642: 151–167. https://doi.org/10.1016/bs.mie.2020.04.070

Gospocic J, Shields EJ, Glastad KM, Lin Y, Penick CA, Yan H, Mikheyev AS, Linksvayer TA, Garcia BA, Berger SL, Liebig J, Reinberg D, Bonasio R. (2017). The neuropeptide corazonin controls social behaviour and caste identity in ants. Cell 10; 170(4): 748–759.e12. https://doi.org/10.1016/j.cell.2017.07.014

Graham RJ. (2018) Epigenetic mechanisms governing behavioural reprogramming in the ant Camponotus floridanus. Publicly Accessible Penn Dissertations. 2883. https://repository.upenn.edu/edissertations/2883

Guo Y, Zhao Z, Li X. (2021). Moderate warming will expand the suitable habitat of *Ophiocordyceps sinensis* and expand the area of *O. sinensis* with high adenosine content. Science of the Total Environment 787: 147605. https://doi.org/10.1016/j.scitotenv.2021.147605

Han Y, van Houte S, Drees GF, van Oers MM, Ros VI. (2015). Parasitic Manipulation of Host Behaviour: Baculovirus SeMNPV EGT Facilitates Tree-Top Disease in Spodoptera exigua Larvae by Extending the Time to Death. Insects 31;6(3): 716-31. https://doi.org/10.3390/insects6030716

Hartze C, Putzier I, Arreola J. (2005). Calcium-activated chloride channels. Annual Review of Physiology 67: 719–758. https://doi.org/10.1146/annurev.physiol.67.032003.154341

Hueffer K, Khatri S, Rideout S, Harris MB, Papke RL, Stokes C, Schulte MK. (2017). Rabies virus modifies host behavior through a snake-toxin like region of its glycoprotein that inhibits neurotransmitter receptors in the CNS. Scientific Reports 7(1):12818. https://doi.org/10.1038/s41598-017-12726-4

Hughes DP, Andersen SB, Hywel-jones NL, Himaman W, Billen J, Boomsma JJ. (2011). Behavioural mechanisms and morphological symptoms of zombie ants dying from fungal infection. BMC Ecology 11(13) 1–10. https://doi.org/10.1186/1472-6785-11-13

Kim D, Ackerman SL. (2011). The UNC5C netrin receptor regulates dorsal mouse hindbrain axons. The Journal of Neuroscience 31(6): 2167–2179. https://doi.org/10.1523/JNEUROSCI.5254-10.2011

Klotz JH, Moss JI. (1996). Oral toxicity of a boric acid – sucrose water bait to Florida carpenter ants (Hymenoptera: Formicidae). Journal of Entomological Science 31: 9–12.

Kopczynski CC, Alton AK, Fechtel K, Kooh PJ, Muskavitch MA. (1988). Delta, a *Drosophila* neurogenic gene, is transcriptionally complex and encodes a protein related to blood coagulation factors and epidermal growth factor of vertebrates. Genes and Development 2(12B): 1723–1735. https://doi.org/10.1101/gad.2.12b.1723

Kumar S, Chen D, Jang C, Nall A, Zheng X, Sehgal A. (2014). An ecdysone-responsive nuclear receptor regulates circadian rhythms in *Drosophila*. Nature Communications 5: 5697. https://doi.org/10.1038/ncomms6697

Lanner JT, Georgiou DK, Joshi AD, Hamilton SL. (2010). Ryanodine Receptors: Structure, expression, molecular details, and function in calcium release. Cold Spring Harbor Perspectives in Biology 2(11): 1003996. https://doi.org/10.1101/cshperspect.a003996

Latchininsky AV, Temreshev II, Childebaev MK, Kolov SV. (2016). Host range and recorded distribution of the fungal pathogen *Entomophaga grylli* (Entomophthoromycota: Entomophthorales) in Kazakhstan. BioONE Complete 25(2): 83–89. https://doi.org/10.1665/034.025.0207

Lavery MN, Murphy CFH, Bowman EK. (2021). View of Zombie ant graveyard dynamics in Gunung Mulu National Park. Reinvention: An International Journal of Undergraduate Research 14(1). https://reinventionjournal.org/article/view/704

Lee K-S, You K-H, Choo J-K, Han Y-M, Yu K. (2004). *Drosophila* short neuropeptide F regulates food intake and body size. Journal of Biological Chemistry 279(49): 50781–50789. https://doi.org/10.1074/jbc.M407842200

Liao Y, Smyth GK and Shi W. (2019). The R package Rsubread is easier, faster, cheaper and better for alignment and quantification of RNA sequencing reads. Nucleic Acids Research 47(8):e47. https://doi.org/10.1093/nar/gkz114

Loreto R, Araújo J, Kepler R, Fleming K, Moreau C, Hughes D. (2018). Evidence for convergent evolution of host parasitic manipulation in response to environmental conditions. Evolution 72(10): 2144–2155. https://doi.org/10.1111/evo.13489

Love MI, Huber W, Anders S. (2014). Moderated estimation of fold change and dispersion for RNA-seq data with DESeq2. Genome Biology 15:550. https://doi.org/10.1186/s13059-014-0550-8

McLaughlin RN Jr, Malik HS. (2017). Genetic conflicts: the usual suspects and beyond. Journal of Experimental Biology 220(1): 6–17. https://doi.org/10.1242/jeb.148148

Mukunda L, Lavista-Llanos S, Hansson BS, Wicher D. (2014). Dimerisation of the *Drosophila* odorant coreceptor Orco. Frontiers in Cellular Neuroscience 28. https://doi.org/10.3389/fncel.2014.00261

Nicholson MJ, Koulman A, Monahan BJ, Pritchard BL, Payne GA, Scott B. (2009). Identification of two aflatrem biosynthesis gene loci in *Aspergillus flavus* and metabolic engineering of *Penicillium paxillin* to elucidate their function. Applied and Environmental Microbiology 75(23): 7469–7481. https://doi.org/10.1128/AEM.02146-08

Nocek B, Mulligan R, Bargassa M, Collart F, Joachimiak A. (2008). Crystal structure of aminopeptidase N from human pathogen Neisseria meningitidis. Proteins 70(1): 273–279. https://doi.org/10.1002/prot.21276

Obulesu M. (2019). Viral Vector Therapeutics Against Alzheimer’s Disease. In: Alzheimer’s Disease Theranostics. Elsevier. https://doi.org/10.1016/C2018-0-00026-2

Pastell M. (2016). CowLog – Cross-platform application for coding behaviours from video. Journal of Open Research Software 4(1): p.e15. https://doi.org/10.5334/jors.113

Pelosi P, Iovinella I, Zhu J, Wang G, Dani F. (2018). Beyond chemoreception: diverse tasks of soluble olfactory proteins in insects. Biological Reviews 93: 184–200. https://doi.org/10.1111/brv.12339

Petch, Trans. Br. Mycol. Soc. 18(1): 53 (1933). Ophiocordyceps clavulata (Schwein.) http://www.indexfungorum.org/Names/NamesRecord.asp?RecordID=251478

Pregitzer P, Greschista M, Breer H, Krieger J. (2014). The sensory neurone membrane protein SNMP1 contributes sensitivity of a pheromone detection system. Insect Molecular Biology 23(6): 733–742. https://doi.org/10.1111/imb.12119

Reichhart N, Schöberl S, Keckeis S, Alfaar AS, Roubeix C, Cordes M, Crespo-Garcia S, Haeckel A, Kociok N, Föckler R, Fels G, Mataruga A, Rauh R, Milenkovic VM, Zühlke K, Klussmann E, Schellenberger E, Strauß O. (2019). Anoctamin-4 is a bona fide Ca^2+^-dependent non-selective cation channel. Scienctific Reports 9: 2257. https://doi.org/10.1038/s41598-018-37287-y

Sakamoto H, Goka K. (2021). Acute toxicity of typical ant control agents to the red imported fire ant, *Solenopsis invicta* (Hymenoptera: Formicidae). Applied Entomology and Zoology 56: 217–224. https://doi.org/10.1007/s13355-021-00728-8

Sánchez MI, Biron DG. (2019). Host Manipulation by Parasites. Frontiers in Ecology and Evolution 7:369. https://doi.org/10.3389/fevo.2019.00369

Selala MI, Daelemans F, Schepens PJC. (2008). Fungal Tremorgens: The mechanism of action of single nitrogen containing toxins – a hypothesis. Drug and Chemical Toxicology 12(3-4): 237–257. https://doi.org/10.3109/01480548908999156

Shakiryanova D, Klose MK, Zhou Y, Gu T, Deitcher DL, Atwood HL, Hewes RS, Levitan ES. (2007). Presynaptic ryanodine receptor-activated calmodulin kinase II increases vesicle mobility and potentiates neuropeptide release. The Journal of Neuroscience 27(29):7799–7806. https://doi.org/10.1523/JNEUROSCI.1879-07.2007

Shinoda T, Itoyama K. (2003). Juvenile hormone acid methyltransferase: A key regulatory enzyme for insect metamorphosis. PNAS 100(21): 11986–11991. https://doi.org/10.1073/pnas.2134232100

Shrestha B, Tanaka E, Hyun MW, Han J-G, Kim CS, Jo JW, Han S-K, Oh J, Sung G-H. (2016). Coleopteran and Lepidopteran hosts of the Entomopathogenic genus *Cordyceps sensu lato*. Journal of Mycology 2016: 7648219. https://doi.org/10.1155/2016/7648219

Sobczak JF, Arruda IDP, Fonseca EO, Rabelo PJQ, Ageu F, Nóbrega dS, Pires JC, Somavilla A. (2020). Manipulation of Wasp (Hymenoptera: Vespidae) Behaviour by the entomopathogenic fungus *Ophiocodyceps humbertii* in the Atlantic forest in Ceará, Brazil. Entomological News 129(1): 98–104. https://doi.org/10.3157/021.129.0115

Steinkraus DC, Hajek AE, Liebherr JK. 2017. Zombie soldier beetles: epizootics in the goldenrod soldier beetle, Chauliognathus 44ensilvanicus (Coleoptera: Cantharidae) caused by Eryniopsis lampyridarum (Entomophthoromycotina: Entomophthoraceae). Journal of Invertebrate Pathology 148: 51–59. https://doi.org/10.1016/j.jip.2017.05.002

Sung G-H, Hywel-Jones NL, Sung J-M, Luangsa-ard JJ, Shrestha B, Spatafora JW. (2007). Phylogenetic classification of *Cordyceps* and the clavicipitaceous fungi. Studies in Mycology 57: 5–59. https://doi.org/10.3114/sim.2007.57.01

Tarailo-Graovac M, Drögemöller BI, Wasserman WW, Ross CJD, van den Ouweland AMW, Darin N, Kollberg G, van Kernebeek CDM, Blomqvist M. (2017). Identification of a large intronic transposal insertion in *SLC17A5* causing sialic acid storage disease. Orphanet Journal of Rare Diseases 12: 28. https://doi.org/10.1186/s13023-017-0584-6

TePaske MR, Gloer JB, Wicklow DT, Dowd PF. (1991). Aflavarin and β-aflatrem: new anti-insectan metabolites from the sclerotia of *Aspergillus flavus*. Journal of Natural Products 55(8): 1080–1086. https://doi.org/10.1021/np50086a008

Tomita H, Cornejo F, Aranda-Pino B, Woodard CL, Rioseco CC, Neel BG, Alvarez AR, Kaplan DR, Miller FD, Cancino G. (2020). The protein tyrosin phosphatase receptor delta regulates developmental neurogenesis. Cell Reports 30(1): 215–228. https://doi.org/10.1016/j.celrep.2019.11.033

Tong EH, Pavey C, O’Handley R, Vyas A. (2021). Behavioral biology of Toxoplasma gondii infection. Parasites and Vectors 14(1):77. https://doi.org/10.1186/s13071-020-04528-x

Torbergsen T. (2002). Rippling muscle disease: a review. Muscle and Nerve 25(11): S103–S107. https://doi.org/10.1002/mus.10156

Trienens M, Rohlfs M. (2011). Experimental evolution of defense against a competitive mold confers reduced sensitivity to fungal toxins but no increased resistance in Drosophila larvae. BMC Evolutionary Biology 11: 206. https://doi.org/10.1186/1471-2148-11-206

Trinh T, Ouellette R, de Bekker C. (2021). Getting lost: the fungal hijacking of ant foraging behaviour in space and time. Animal Behaviour 181:165–184. https://doi.or/10.1016/j.anbehav.2021.09.003

Valdes JJ, Cameron JE, Cole RJ. (1985). Aflatrem: A tremorgenic mycotoxin with acute neurotoxic effects. Environmental Health Perspectives 62: 459–463. https://doi.org/10.1289/ehp.8562459

Wells L, Edwards KA, Bernstein SI. (1996). Myosin heavy chain isoforms regulate muscle function but not myofibril assembly. The EMBO Journal 15(17): 4454–4459. https://doi.org/10.1002/j.1460-2075.1996.tb00822.x

Werkhoven Z, Rohrsen C, Qin C, Brembs B, de Bivort B. (2019). MARGO (Massively Automated Real-time GUI for object-tracking), a platform for high-throughput ethology. PLoS one 14(11): e0224243. https://doi.org/10.1371/journal.pone.0224243

Wesołowska W, Wesołowski T. (2013). Do *Leucochloridium* sporocysts manipulate the behaviour of their snail hosts? Journal of Zoology 292(3): 151–155. https://doi.org/10.1111/jzo.12094

Will I, Linehan S, Jenkins DG, de Bekker C. (Under review). Natural history and ecological effects on the establishment and fate of Florida carpenter ant cadavers infected by the parasitic manipulation Ophiocordyceps camponoti-floridani

Will I, Das B, Trinh T, Brachmann A, Ohm RA, de Bekker C. 2020. Genetic underpinnings of host manipulation by *Ophiocordyceps* as revealed by comparative transcriptomics. G3 10(7): 2275–2296. https://doi.org/10.1534/g3.120.401290

Wnuk A, Kostowski W, Korczyńska J, Szczuka A, Symonowicz B, Bieńkowski P, Mierzejewski P, Godzińska EJ. (2014). Brian GABA and glutamate levels in workers of two ant species (Hymenoptera: Formicidae): interspecific differences and effects of queen presence/absence. Insect Science 5: 647–658. https://doi.org/10.1111/1744-7917.12076

Xing X, Yan M, Pang H, Wu F, Wang J, Sheng S. (2021). Cytochrome P450s Are Essential for Insecticide Tolerance in the Endoparasitoid Wasp Meteorus pulchricornis (Hymenoptera: Braconidae). Insects 12(7):651. https://doi.org/10.3390/insects12070651

Yao Y, Peter AB, Baue R, Sigel E. (1989). The tremorigen aflatrem is a positive allosteric modulator of the gamma-aminobutyric acidA receptor channel expressed in Xenopus oocytes. Molecular Pharmacology 35(3): 319–323. https://molpharm.aspetjournals.org/content/35/3/319.long

Zalewska M, Kochman A, Estève JP, Lopez F, Chaoui K, Susini C, Ozyhar A, Kochman M. (2009). Juvenile hormone binding protein traffic - Interaction with ATP synthase and lipid transfer proteins. Ciochimica et Biophysica Acta 1788(9):1695–705. https://doi.org/10.1016/j.bbamem.2009.04.022

Zhang Y, Featherstone D, Davis W, Rushton E, Broadie K. (2000). *Drosophila* D-Titin is required for myoblast fusion and skeletal muscle striation. Journal of Cell Science 113: 3103–3115. https://doi.org/10.1242/jcs.113.17.3103

